# Patterns of subcortical tissue iron levels across the lifespan measured with functional MRI

**DOI:** 10.64898/2026.09.09.750391

**Authors:** Attakias T. Mertens, Derek J. Pavelka, Lauren Foley, Katrina Myers, Callum Goldsmith, Tatiana Wolfe, Jacob J. Oleson, Gaelle E. Doucet

**Affiliations:** Institute for Human Neuroscience, Boys Town National Research Hospital, Boys Town, NE, USA; Brain Imaging Research Center, Psychiatric Research Institute, Winthrop P. Rockefeller Cancer Institute, College of Medicine, University of Arkansas for Medical Sciences, Little Rock, AR, USA; University of Iowa, Iowa City, IA, USA; Center for Pediatric Brain Health, Boys Town National Research Hospital, Boys Town, NE, USA; Department of Pharmacology and Neuroscience, Creighton University School of Medicine, Omaha, NE, USA

**Keywords:** subcortical iron, typical development, healthy aging, lifespan, functional magnetic resonance imaging

## Abstract

Iron is essential for neurophysiological health. Prior work has shown that subcortical tissue iron levels are the highest among all brain regions but also change over the lifespan. Recent functional MRI (fMRI) studies have provided evidence that an fMRI-derived iron-sensitive signal may index relative iron concentration in subcortical regions. In this context, we aimed to characterize the lifespan trajectory of this fMRI-derived iron-sensitive signal (relative iron quantification) in 10 subcortical regions and 14 thalamic nuclei from 881 healthy participants aged 12 to 88 years, using resting-state fMRI. Regressions were run with age as a predictor, separately for males (N = 429) and females (N = 452). The main results show non-linear effects of age, with an overall increase in iron levels across most subcortical regions within each sex. Six of the 14 thalamic nuclei in male participants had similar slopes of their lifespan iron levels, showing some homogeneity. The thalamic nuclei of female participants showed less agreement in slopes. This study provides new insight into the typical trajectory of iron levels across all major subcortical regions throughout the lifespan.

## 1. Introduction

Intracellular, non-heme iron is the most prevalent metal in the brain (Hect et al., 2018). Previous postmortem studies established that subcortical regions exhibit higher iron levels than the cortex, and that these levels vary by region and lifespan stage (Hallgren & Sourander, 1958; Langkammer et al., 2010, 2012; Ramos et al., 2014). In early life, iron promotes neural health and development. It is required for the synthesis of many neurotransmitters, including dopamine (DA), norepinephrine, serotonin, GABA, and L-glutamate (Hidalgo & Núñez, 2007; Kulaszyńska et al., 2024; Piñero & Connor, 2000). Iron is also prevalent in oligodendrocytes, where it aids in the creation and maintenance of myelin (McCann et al., 2020; Piñero & Connor, 2000; Todorich et al., 2009). In addition to its neurophysiological contributions, higher levels of regional brain iron are associated with better cognitive performance across domains in typically developing children, adolescents, and young adults (Hect et al., 2018; Larsen et al., 2020). In contrast, certain developmental disorders show differences in subcortical iron concentration, such as children with attention-deficit/hyperactivity disorder (ADHD) who generally show lower levels compared to typically developing children, especially within the thalamus (Adisetiyo et al., 2019; Cortese et al., 2012; Hasaneen et al., 2017). These findings remain contentious, however, given other reports showing no differences between these groups (Cascone et al., 2023; Schulze et al., 2025). Aside from this, higher caudate tissue iron is associated with lower performance on a verbal memory task in children with developmental language disorder (DLD) but not in typically developing children (Doucet et al., 2025).

In contrast to early life development, higher subcortical tissue iron levels in older adults tend to have more detrimental effects on neural health. While brain tissue iron levels are reported to increase throughout healthy aging, elevated levels are associated with poorer myelin integrity, increased apoptosis, grey matter atrophy, inflammation, metabolic dysfunction and poorer cognitive performance (Andersen et al., 2014; Daugherty & Raz, 2013; Dixon & Stockwell, 2014; Madden & Merenstein, 2023; Spence et al., 2020). Abnormal levels are also reported in various diseases/disorders, such as the higher thalamic tissue iron found in adults with schizophrenia (Sonnenschein et al., 2022). Furthermore, patients with Parkinson’s disease (PD) show higher substantia nigra (SN) tissue iron, particularly in the pars compacta (SNpc), compared to healthy individuals (López-Aguirre et al., 2025; Wang et al., 2022). Meanwhile, higher hippocampal tissue iron is associated with mild cognitive impairment (MCI) and Alzheimer’s disease (AD; Kagerer et al., 2025; Mieling et al., 2025; Zeineh et al., 2015). This suggests that tissue iron accumulation may be a potential indicator of AD development from MCI. Along with this, higher inferior temporal gyrus tissue iron is associated with increased tau accumulation and lower global cognitive functioning in AD (Spotorno et al., 2020).

Advances in magnetic resonance imaging (MRI) techniques and technology present the potential for non-invasive, in-vivo measurement of brain iron. One such approach obtains a relative, iron-sensitive fMRI signal by normalizing and averaging T2*-weighted images over time, which are extracted from echo-planar imaging sequences typically used in the context of fMRI. This approach has been implemented previously to investigate iron concentration in relation to cognitive processes (Larsen et al., 2020; Parr et al., 2022), typical development (Larsen & Luna, 2015), children with DLD (Doucet et al., 2025), children with ADHD (Cascone et al., 2023), substance usage during adolescence (Flannery et al., 2025; Parr et al., 2026), depression (Price et al., 2021), and schizophrenia (Sonnenschein et al., 2022).

Overall, the usage of fMRI offers the potential for more widespread investigation of tissue iron levels within the brain, especially considering resting-state fMRI (rs-fMRI) is among the most commonly collected sequences in research MRI protocols, in contrast to other MRI sequences which are not consistently collected (e.g., quantitative MRI, quantitative susceptibility mapping). For example, the Human Connectome Project (HCP; Elam et al., 2021; Glasser et al., 2013, 2016), Autism Brain Imaging Data Exchange (ABIDE; Di Martino et al., 2014, 2017), and Parkinson’s Progression Markers Initiative (PPMI; Marek et al., 2011) all collect rs-fMRI.

While multiple studies have found evidence that subcortical tissue iron levels generally increase throughout life, the exact slope of this increase remains unclear. Prior studies have differed in the regions of interest (ROIs), age range, and types of slopes investigated. For example, one study used five ROIs from participants aged 10-70 years, and assessed three types of slopes, including linear, quadratic, and exponential curves (Li et al., 2023). In contrast, another study investigated eight ROIs’ tissue iron levels, but only in adults aged 20-70 for linear and quadratic slopes (Persson et al., 2015). One of the largest lifespan studies to date included 498 participants aged 5-90 years (Treit et al., 2021). This study assessed a wider range of non-linear slopes, but only in four ROIs. Altogether, there are still several subcortical regions whose typical lifespan tissue iron trajectories have yet to be investigated, including the hippocampus, nucleus accumbens or red nucleus, among others.

Currently, there is a lack of lifespan studies on subcortical tissue iron utilizing fMRI data. In this context, this study aims to provide a more complete picture of the lifespan trajectory of tissue iron: 1) in a larger age range, from childhood to late adulthood, 2) using a larger set of subcortical regions, than previous studies, and 3) testing both linear and non-linear slopes to better characterize these age-related changes. For this, we used rs-fMRI data from 881 healthy participants aged 12-88 years. All analyses were conducted separately within each sex to determine any potential sex-specific age trajectories.

## 2. Methods

### 2.1 Participants

Data from two data sets was used in this study. In total, the sample included 881 healthy participants aged 12 to 88 years (51% female, mean age (SD) = 47.56 (21.78), Figure 1).

**Figure 1:**
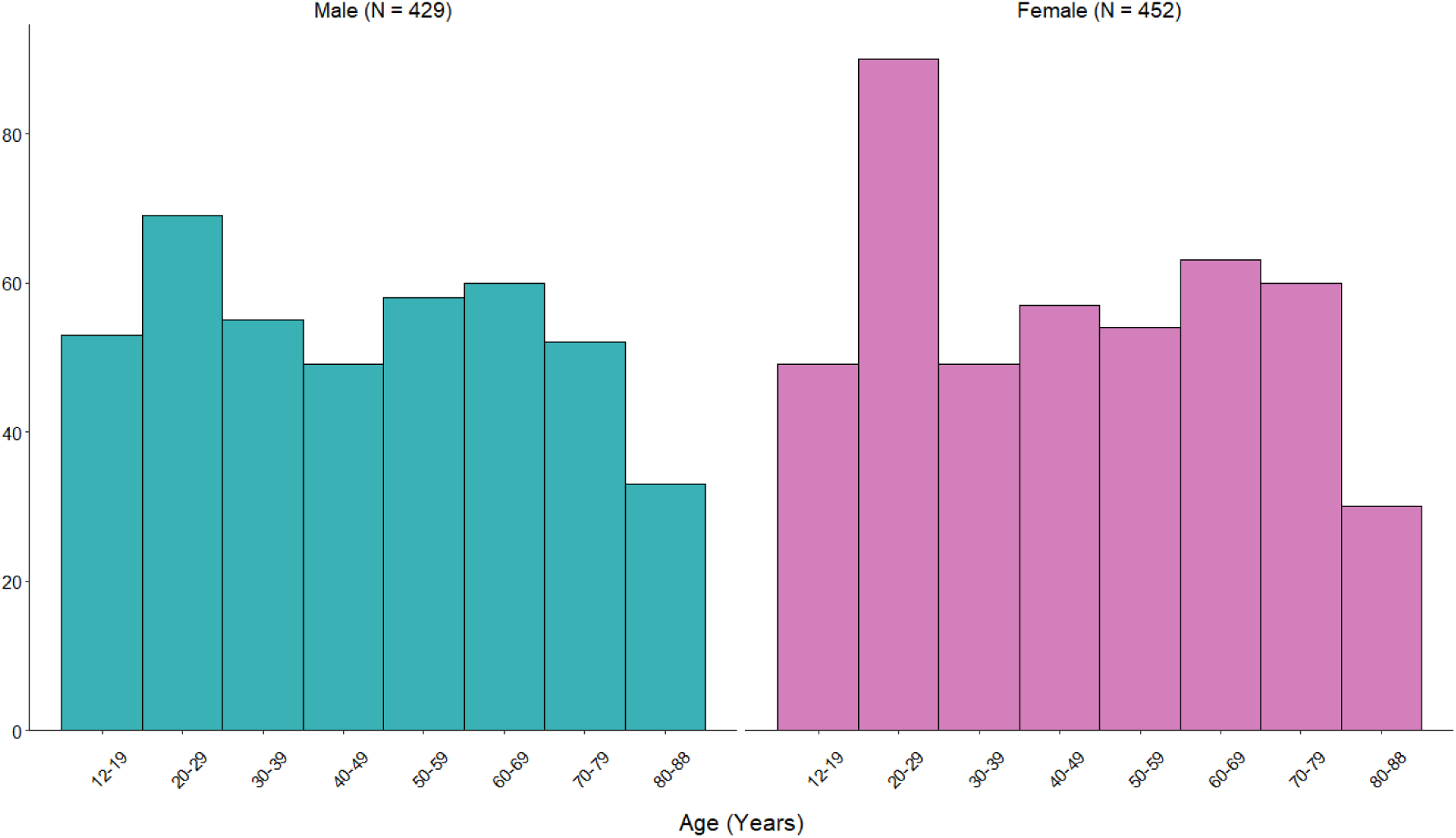
Age Histograms

The first and largest dataset was based on the publicly available the Cambridge Centre for Aging and Neuroscience study (CamCAN). This study used an epidemiologically informed recruitment framework and is available upon request (https://www.cam-can.org/; Shafto et al., 2014; Taylor et al., 2017). From this cohort, we selected participants who had completed an rs-fMRI scan and had demographic data. This included 644 healthy participants aged 18 to 88 (mean age (SD) = 54.89 (18.55) years, 51% female). This study was approved by the Cambridgeshire 2 Research Ethics Committee. Participants agreed to participate in the research by providing written consent. Details on participant recruitment are provided in Taylor et al. (2017).

The second dataset was locally collected and included a total of 237 typically developing and healthy participants ages 12 to 81 (mean age (SD) = 27.63 (16.91) years, 52% female). The main reason to add this dataset was to have brain iron information for adolescents (70 were 18 years old or under) as they were not available in the CamCAN sample (which is focused on adulthood). Further, this dataset also included adults, which can provide evidence of reliability and generalizability of the findings across independent datasets that used different scanners and MRI sequences. Overall, having these two datasets provided a more uniform age distribution for each decade, from 12 to 88 years old (Figure 1). The general exclusion criteria included individuals who had a diagnosis of a psychiatric or neurologic disorder, had a history of drug use or did not meet safety criteria for the MRI scan (metal implants, pregnancy, etc.). Research was approved by the institutional review board (IRB) from the local committee. Participants and/or their legal guardian consented to participate in the research, while the children provided written assent.

### 2.2 MRI Acquisition and Preprocessing

MRI data were collected through two research sites, though both used a 3 T Siemens Prisma scanner. Acquisition parameters of structural and rs-fMRI images differed by site.

Acquisition parameters used in the CamCAN sample include for the structural image (T1-weighted, MPRAGE) were: Repetition Time (TR) = 2250ms, Echo Time (TE) = 2.99ms, Inversion Time (TI) = 900ms, 9 degree-flip angle, Field of View (FOV) = 256 x 240 x 192mm, voxel size = 1 x 1 x 1mm^3^, acquisition time = 4m and 32s, and GRAPPA factor 2. For the rs-fMRI scans, the T2* weighted EPI sequence included: TR = 1970 ms, TE = 30ms, flip angle = 78 degrees, FOV = 192 x 192 mm, flip angle = 78 degrees, volumes = 261, number of axial slices = 32, acquisition time = 8m and 40 s, and voxel size = 3 x 3 x 4.4 mm.

For our local sample (*n* = 238), parameters for the structural image (T1-weighted, 3D magnetization-prepared rapid gradient-echo (MPRAGE)) were: TR = 2400ms, TE = 2.22ms, FOV= 256 × 256mm, matrix size: 320 × 320, 0.8mm^3^ isotropic resolution, TI = 1000 ms, 8 degree-flip angle, bandwidth = 220 Hz/Pixel, echo spacing = 7.5ms, in-plane acceleration GRAPPA (GeneRalized Autocalibrating Partial Parallel Acquisition) factor 2. For the rs-fMRI scans, the T2* EPI sequence had the following acquisition parameters: TR = 800ms, TE = 37ms, voxel size = 2 × 2 × 2mm^3^, echo spacing = 0.58ms, total volumes = 700, bandwidth = 2290 Hz/Pixel, number of axial slices = 72, multi-band acceleration factor = 8.

Both rs-fMRI datasets were similarly preprocessed using *fMRIprep* v.23.0.2 using the default pipeline, which included slice-time correction, realignment, coregistration, and normalization.

### 2.3 Regions of Interest (ROIs)

All ROIs used in this study were extracted using the Automated Anatomical Labeling (AAL3) structural atlas (Rolls et al., 2020). The main 10 ROIs chosen were focused on subcortical regions previously shown to have high brain iron content (Acosta-Cabronero et al., 2016; Cascone et al., 2023; Hect et al., 2018; Larsen et al., 2020; Petok et al., 2024) or have altered iron concentrations related to a disorder (Acosta-Cabronero et al., 2013; Adisetiyo et al., 2019; Ahmadi et al., 2020), including the caudate, hippocampus, nucleus accumbens (NA), pallidum, putamen, red nucleus (RN), substantia nigra pars compacta (SNpc), substantia nigra pars reticulata (SNpr), thalamus, and ventral tegmental area (VTA).

Given the heterogeneity of previous findings in regard to thalamic tissue iron trajectory across the lifespan (Acosta-Cabronero et al., 2016; Bilgic et al., 2012; Haacke et al., 2010; Treit et al., 2021), we chose to further investigate 14 major thalamic nuclei. All of these can be found in Table 1 listed with their abbreviations. The reuniens nucleus was excluded, due to its small volume (8mm^3^; Rolls et al., 2020).

**Table 1:**
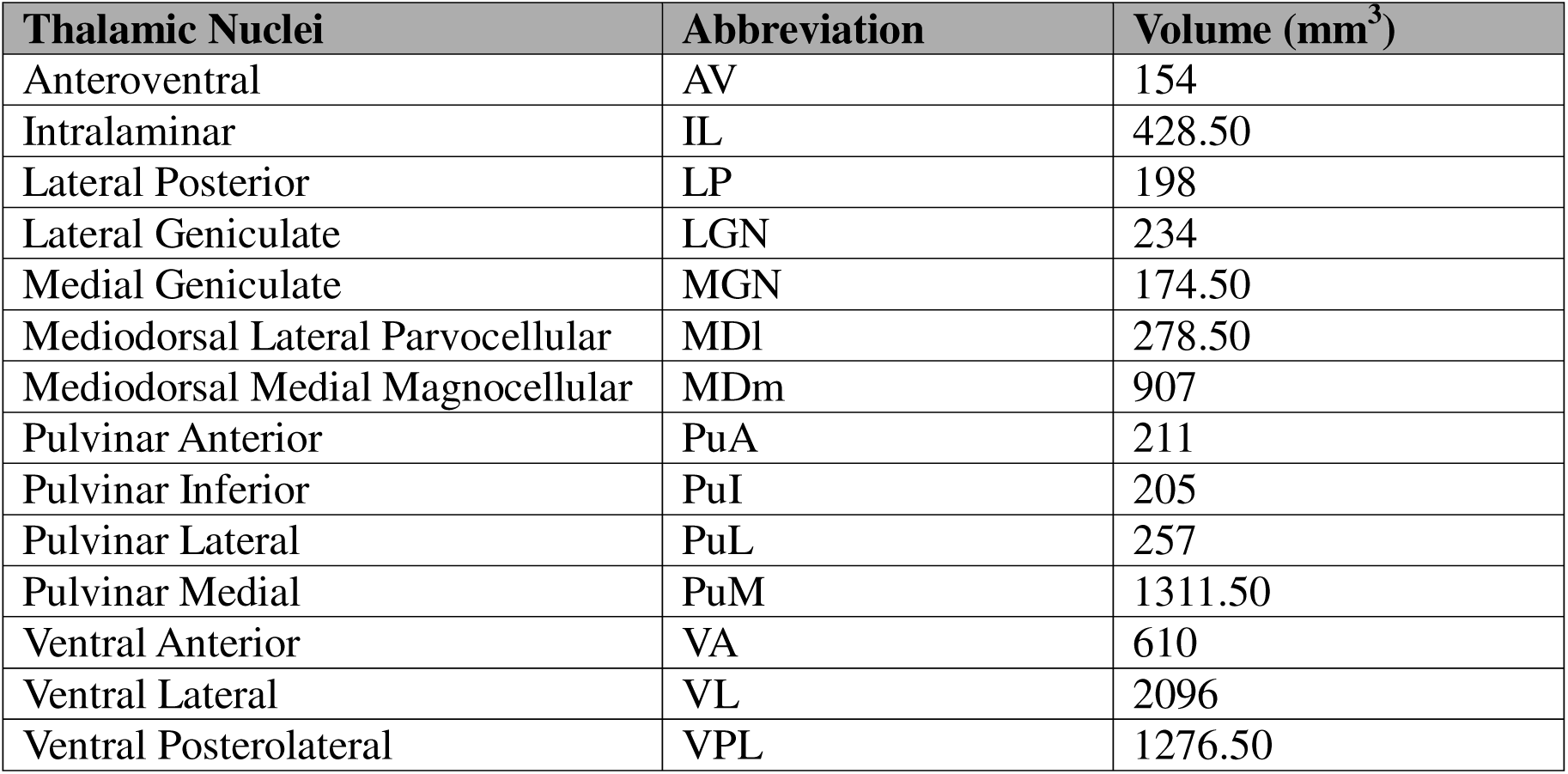
Thalamic Nuclei used in the study with their abbreviations.

| Thalamic Nuclei | Abbreviation | Volume (mm<sup>3</sup>) |
| --- | --- | --- |
| Anteroventral | AV | 154 |
| Intralaminar | IL | 428.50 |
| Lateral Posterior | LP | 198 |
| Lateral Geniculate | LGN | 234 |
| Medial Geniculate | MGN | 174.50 |
| Mediodorsal Lateral Parvocellular | MDl | 278.50 |
| Mediodorsal Medial Magnocellular | MDm | 907 |
| Pulvinar Anterior | PuA | 211 |
| Pulvinar Inferior | PuI | 205 |
| Pulvinar Lateral | PuL | 257 |
| Pulvinar Medial | PuM | 1311.50 |
| Ventral Anterior | VA | 610 |
| Ventral Lateral | VL | 2096 |
| Ventral Posterolateral | VPL | 1276.50 |

As previous studies have reported no hemispheric differences in these regions (Acosta-Cabronero et al., 2016; Sonnenschein et al., 2022), the right and left regions were averaged for each of the total 24 ROIs, in line with previous studies (Burgetova et al., 2021; Larsen et al., 2020; Treit et al., 2021).

### 2.4. Brain Iron Quantification

The brain iron quantification process in this study followed the approach used in prior rs-fMRI works (Doucet et al., 2025; Larsen & Luna, 2015; Parr et al., 2022, 2026; Peterson et al., 2018). Briefly, each volume of the preprocessed rs-fMRI was normalized to its own mean within a brain mask created during the preprocessing step. Volumes with a framewise displacement *>* 0.3 were excluded. The T2*-weighted (T2*w) signal originates from magnetic susceptibility, and it reflects contributions from both paramagnetic (e.g., iron) and diamagnetic (e.g., myelin) tissue sources; it is therefore iron-sensitive, but not iron-specific. However, it has been found that T2*w is more sensitive to paramagnetic iron load than diamagnetic (e.g., myelin) tissue sources (Larsen & Luna, 2015). Next, the median across all included normalized volumes was computed for each brain voxel. This process resulted in one normalized T2*w (nT2*w) image for each participant. Mean values of this signal were then computed within each ROI. Next, these values were inverted (1/nT2*w) to provide an fMRI-derived iron-sensitive measure that reflects a relative, rather than absolute, iron R2* map (rR2*). In this case, a higher rR2* signal is expected to covary with and be sensitive to higher iron content, while a lower rR2* signal is expected to reflect lower iron content. Given that differences in MRI scanning parameters, software, and scanner locations can result in signal variation, values for rR2* were site corrected using ComBat Harmonization v1.0.1 (Fortin et al., 2017, 2018) for MATLAB (2023a). Harmonized outputs for participants from each site within the overlapping age range (between 50 and 80 years old) showed no significant site effect after harmonization for all ROIs (range *p* = .661 - .941). Therefore, harmonized values for rR2* were used from this point. Overall regional averages aligned with patterns found in previous studies (Figure 2; Aquino et al., 2009; Bilgic et al., 2012; Ghadery et al., 2015; Haacke et al., 2010; Hallgren & Sourander, 1958; Hect et al., 2018; Langkammer et al., 2010, 2012; Larsen et al., 2020; Péran et al., 2009; Persson et al., 2015). Python 3 (v3.11.13) toolboxes nibabel (v5.3.2; Brett et al., 2025) and numpy (v2.3.1; Harris et al., 2020) were used.

**Figure 2:**
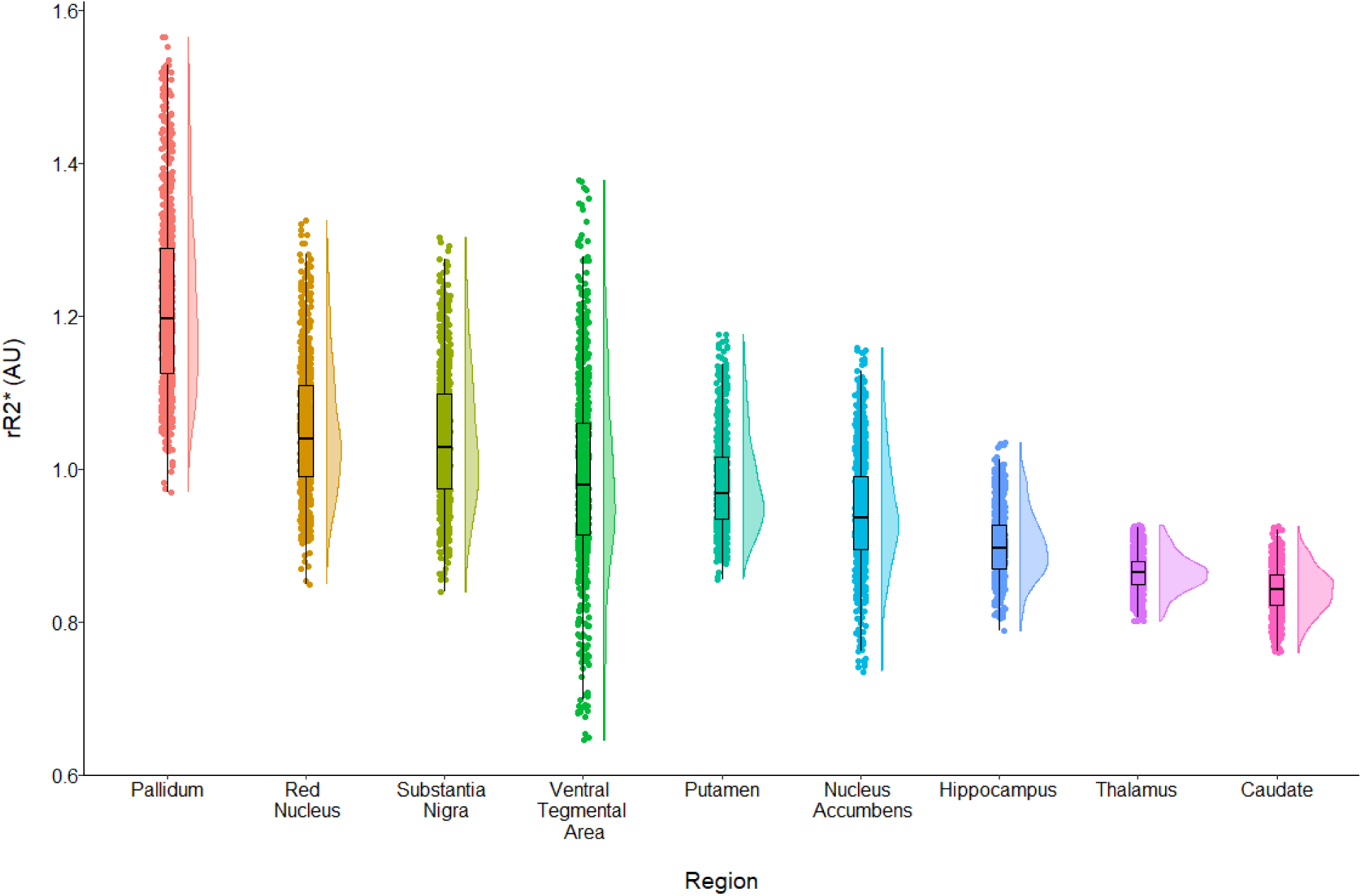
Raincloud Plots for Main Regions of Interest’s rR2* *Note*. For each region of interest: 1) the right side shows the distribution of rR2* values, 2) the left side shows dots representing individual participant values and boxplots show regional median values, along with the first and third quartiles.

### 2.5. Statistical Analysis

Our main goal was to map brain iron content in each subcortical region across the lifespan. For this, we conducted a series of regression analyses with age as the main predictor. Given that previous studies have shown age to have a nonlinear relationship with brain iron content, we ran five regression models to test different curves in each ROI: linear, quadratic, cubic, logarithmic, and s-curve. Outliers for each ROI were identified using interquartile range. Mean-centered age was used in models testing multiple regression, including the quadratic and cubic models. Models with severe heteroskedasticity were rerun using weighted least squares regression. The best model was identified as having the lowest Akaike Information Criterion (AIC). All statistics were conducted with R v.4.4.3 in RStudio.

Previous findings on the differences between sexes across different age ranges for iron content in subcortical regions are mixed (Hect et al., 2018; Larsen et al., 2020; G. Li et al., 2023; Persson et al., 2015; Peterson et al., 2018). Thus, we chose to run all regression analyses separately for female and male participants to determine their independent trajectories. Multiple comparisons were corrected for with a false discovery rate (FDR) correction (Benjamini & Hochberg, 1995) for the analyses of the main 10 ROIs and the 14 thalamic nuclei, separately.

### 2.6. Supplementary Analyses

The link between qMRI and post-mortem studies has been demonstrated using the same rank-order convergent validity approach -the accepted validation standard across all MRI iron methods-which consists of correlating in vivo regional MRI values against the established post-mortem regional iron hierarchy (Bilgic et al., 2012; Feng et al., 2018; Hallgren & Sourander, 1958; Langkammer et al., 2010, 2012; Pfefferbaum et al., 2009; Spincemaille et al., 2019; Xu et al., 2008; Yan et al., 2012). Pfefferbaum et al. (2009), specifically, applied this strategy to validate two QSM approaches and found the same rank ordering across structures. Therefore, we compared the averaged regional iron concentrations reported in previous post-mortem work (Hallgren & Sourander, 1958) with those of the current rR2*. These values were calculated within the current caudate, pallidum, putamen, RN, the average of the SN_pc and SN_pr (SN), and thalamus of participants aged 30 and above (*N* = 395) to align with the age range reported by Hallgren and Sourander (1958). We confirmed an extremely high correlation (r=.92, p=.009), which is similar or higher than the relationship previously reported between qMRI and post-mortem data (Langkammer et al., 2010, 2012; Pfefferbaum et al., 2009; Xu et al., 2008).

## 3. Results

### 3.1 Regression analyses in the 10 main subcortical regions

All model results for each ROI are provided in the Supplementary Materials (Supplementary Table 1). The models including the best fit, or lowest AIC (Supplementary Table 2), for each ROI, are reported in Table 2 and described below in accordance with the general trend in the slope they have for the association between age and rR2* (Figure 3).

**Table 2:**
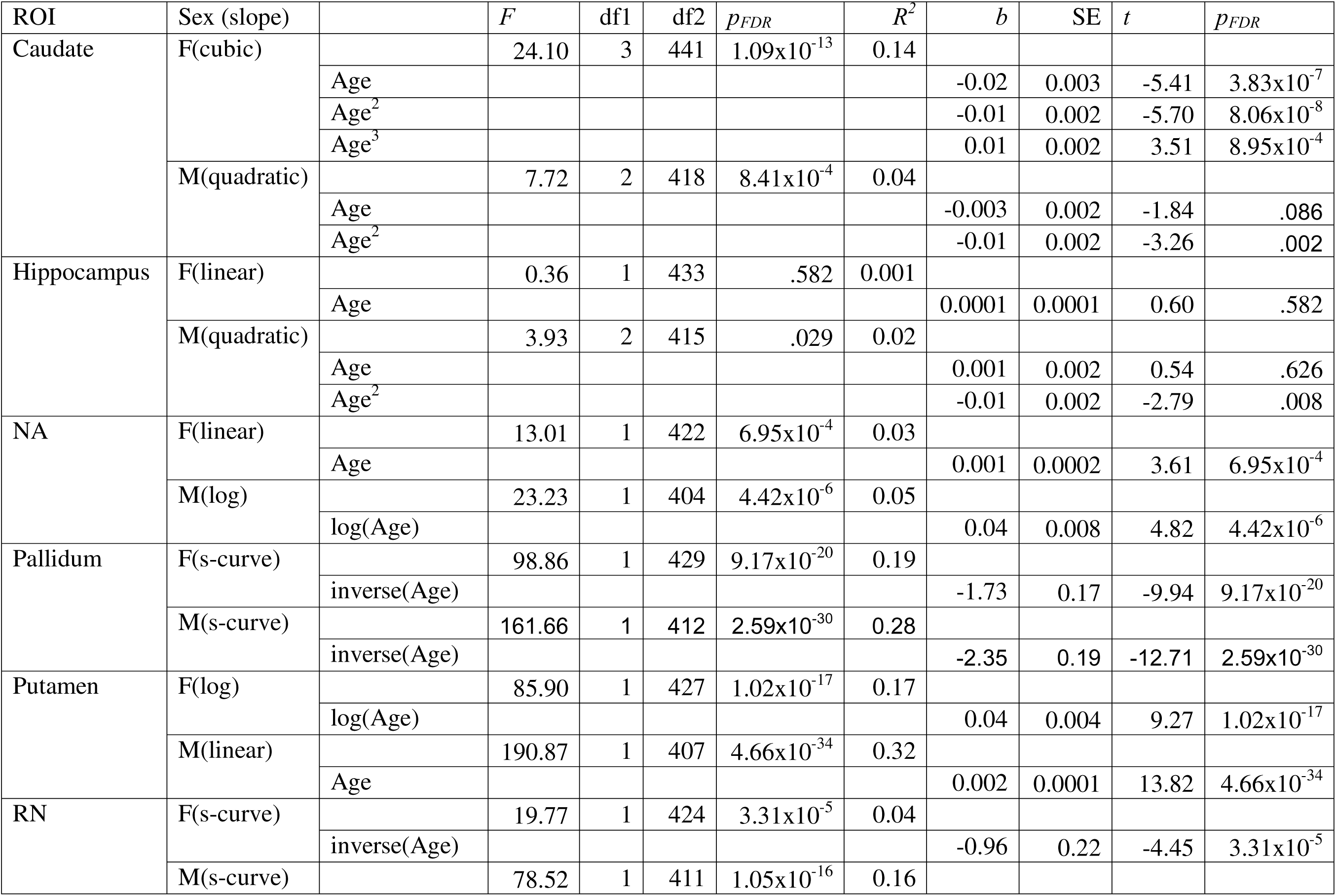
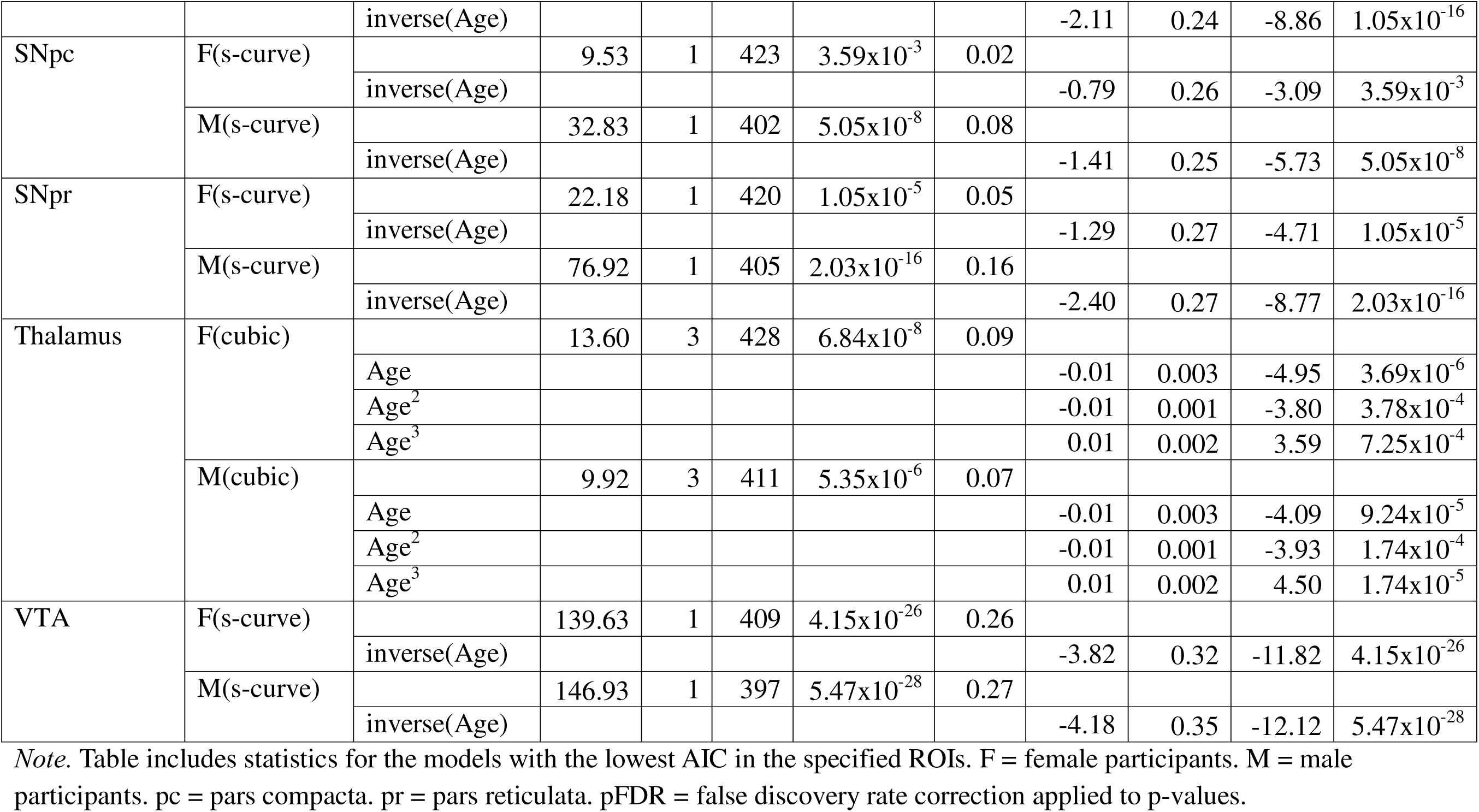
Best Fit Models for ROIs with General Increasing Slopes in rR2* with Age.

**Figure 3:**
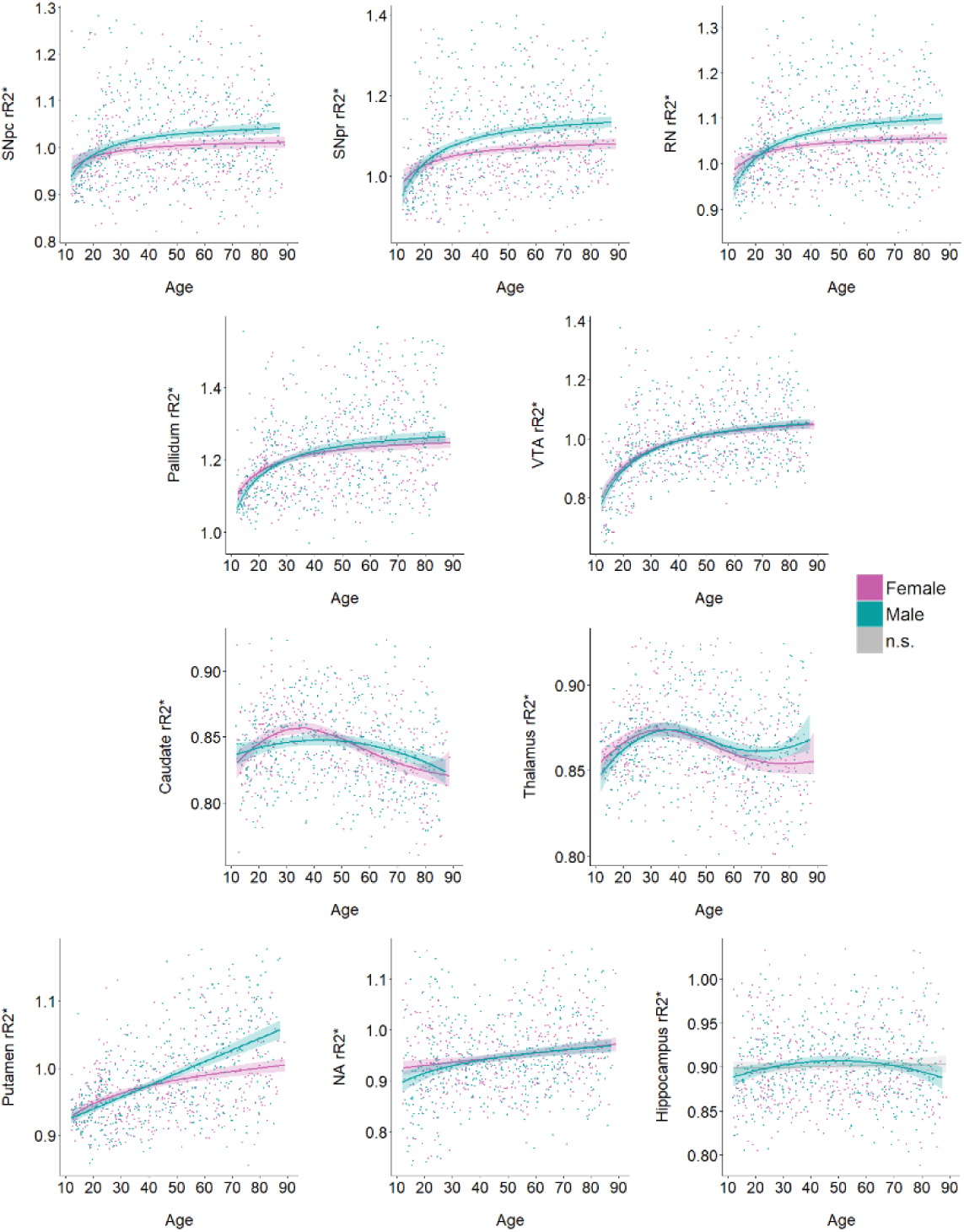
Best Fit Curves for Main Regions of Interest’s rR2* *Note*. Figure shows significant curves of rR2* in the specified regions across age in female (purple) and male (blue) participants. Both sexes are graphed together for visualization, though models for each sex were run separately. pc = pars compacta. pr = pars reticulata. RN = red nucleus. VTA = ventral tegmental area. NA = nucleus accumbens. n.s. = not significant.

Half of the main ROIs’ rR2* showed steeper, increasing slopes primarily during early development (∼ <30 years), followed by positive, though more flat slopes throughout later life. This included female and male participants’ pallidum, RN, SNpc, SNpr, and VTA (s-curve). The thalamus showed a cubic best-fit slope for both sexes, with increasing rR2* (higher tissue iron levels) in early development (< ∼30 years), followed by a decrease throughout midlife and into older age (30s to 70s). Female participants showed a highly similar slope for the caudate (cubic) compared to the thalamus, while male participants differed (quadratic). For the caudate, male participants showed an initial increase in rR2* (higher tissue iron levels) until middle adulthood (∼ <50 years), followed by a decrease. A general increase in rR2* (higher tissue iron levels) was observed in both sexes putamen, with female participants (log) showing a greater rise in early life (∼ <30 years) followed by a more gradual increase thereafter and male participants (linear) showing a consistent rise across the lifespan. Similarly, an overall positive slope was found for both sexes NA, however; female participants (linear) showed a gradual increase across age for this region, while male participants (log) had a steeper, though moderate, increase in early life (∼ <30 years), compared to their gradual increase that followed. Lastly, for the hippocampus, male participants (quad) showed a slight increase until midlife (< ∼50 years), followed by a slight decrease, while female participants did not have a significant age association.

### 3.2 Regression analyses in the 14 main thalamic nuclei

The thalamic nuclei’s rR2* varied in their slopes of best fit and displayed three primary patterns (Table 3; Figure 4). The first included cubic trends that were somewhat similar to the overall thalamic slope. The MDl, MDm, and PuI showed this pattern for both sexes. This same cubic slope was also found in female participants’ IL and male participants’ AV, PuL, and VA. Additionally, male participants also had a cubic slope in their LP, though it differed from the overall thalamus slope in that it showed a plateau in late life. The second pattern included a quadratic slope, seen in female participants’ AV and VA, where iron levels increased until midlife (< ∼50 years) and then decreased. In male participants, a quadratic slope was found in the LGN and PuA, where iron levels increased until midlife (< ∼60 years) and then decreased, and in the MGN and PuM which similarly increased until midlife (< ∼50 years) and then decreased.

**Table 3:**
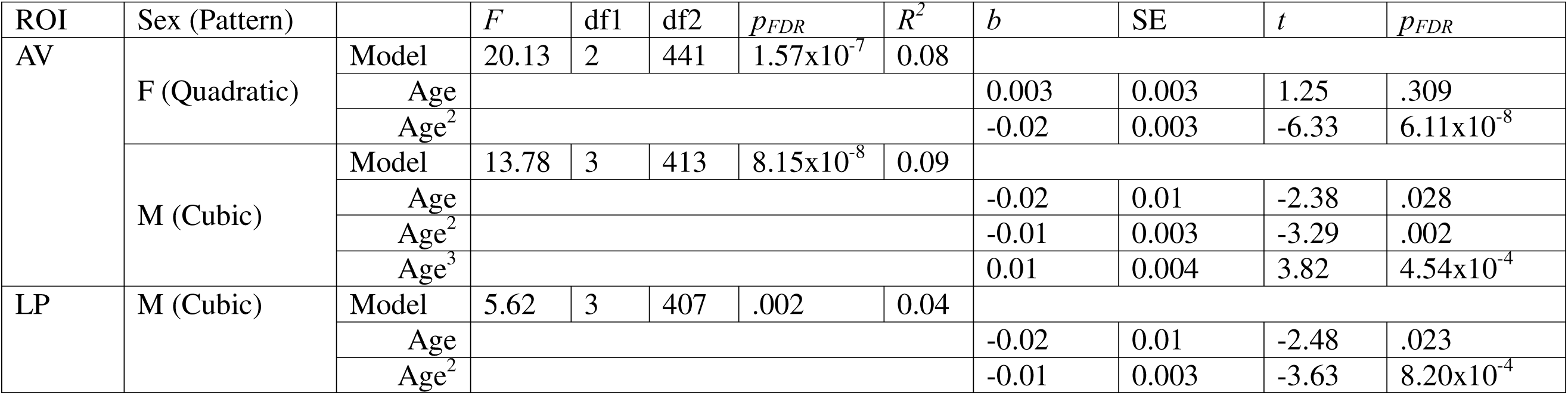
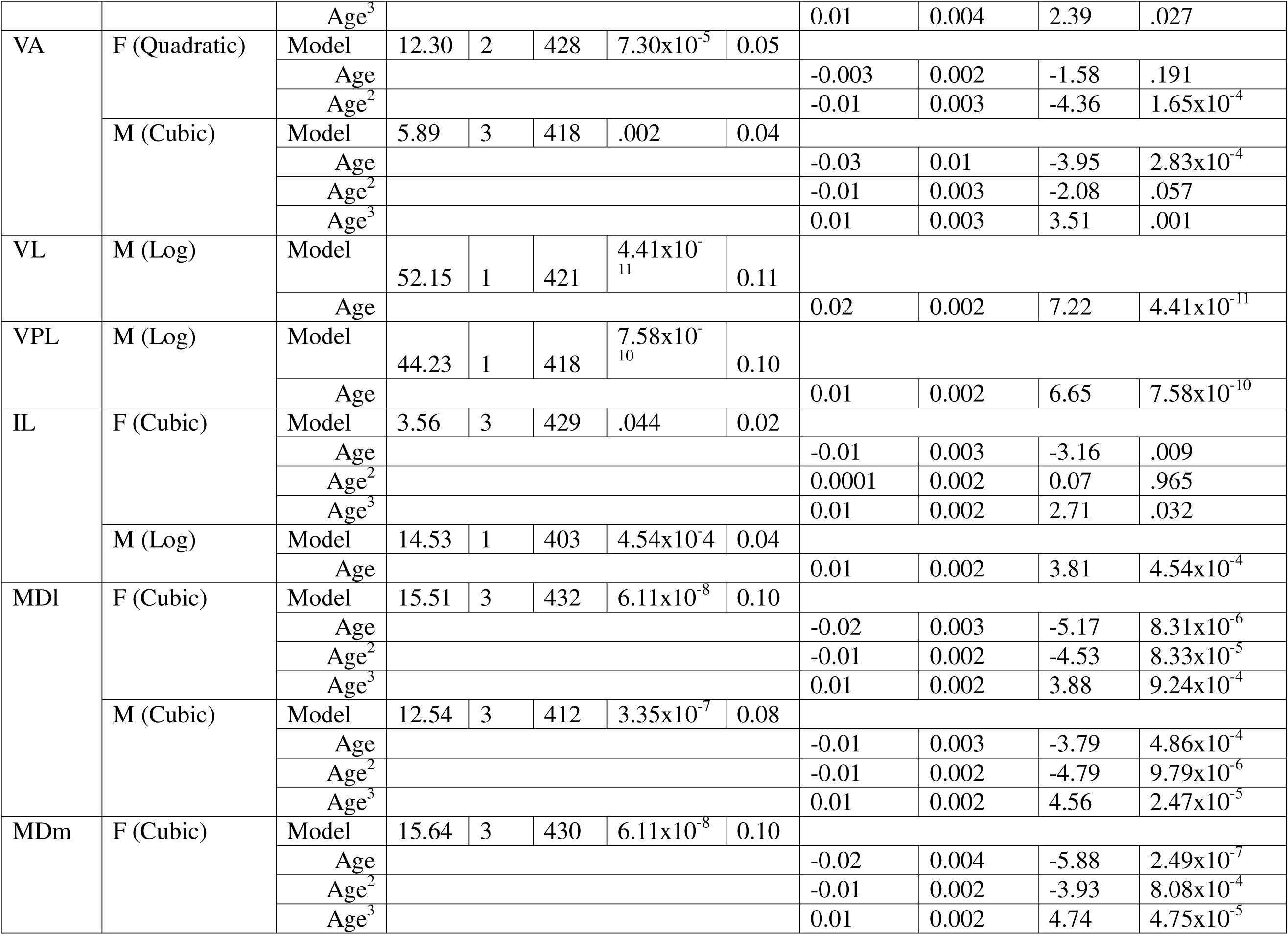
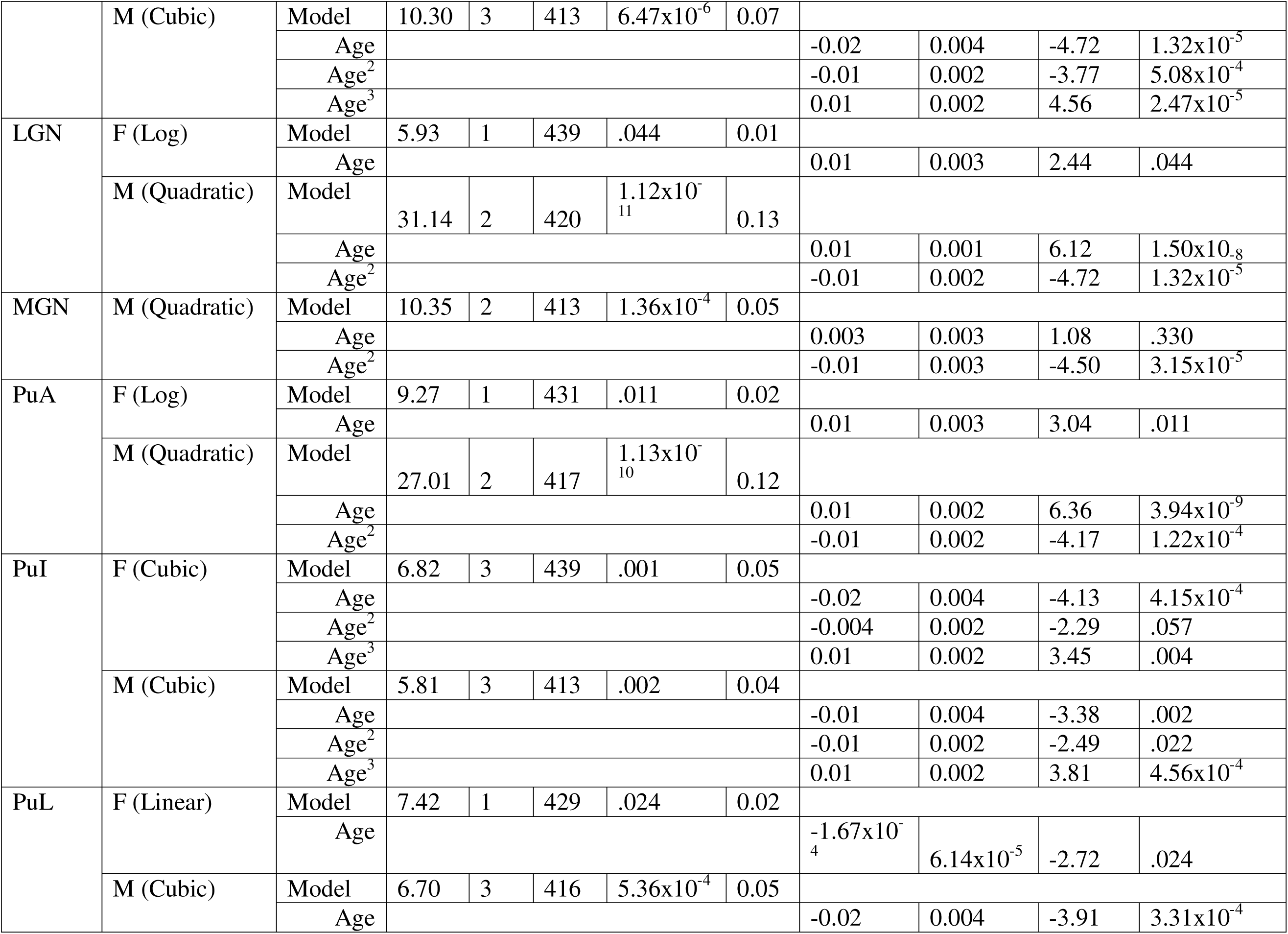
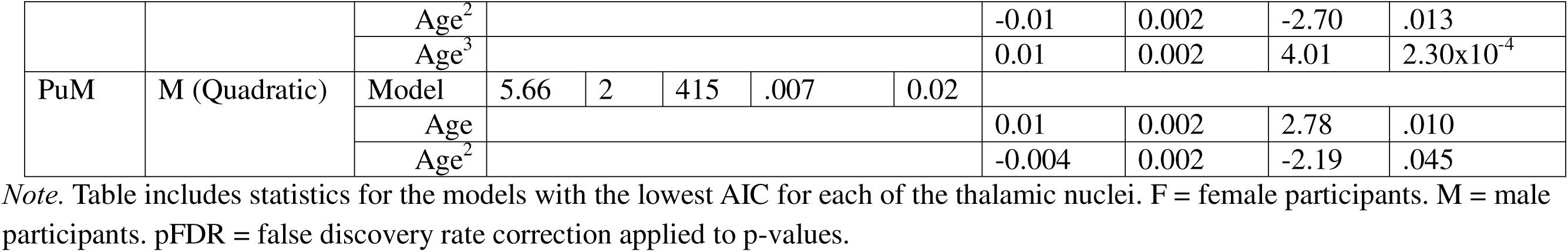
Best Fit Models for Thalamic Nuclei’s rR2*.

**Figure 4:**
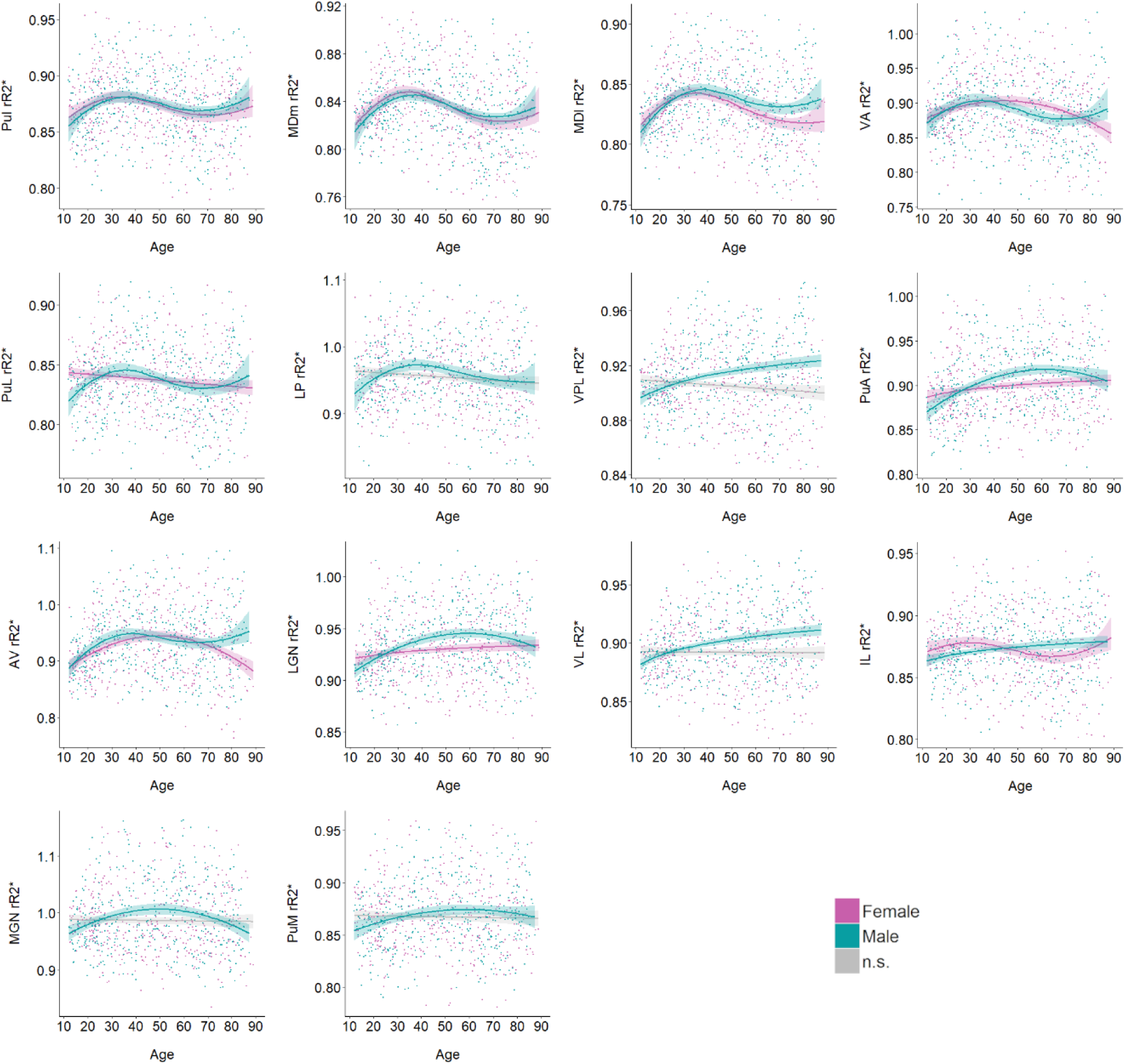
Best Fit Curves for Thalamic Nuclei’s rR2* *Note.* Figure shows significant curves of iron content in the specified thalamic nuclei across age in female and male participants. Both sexes are graphed together for visualization, though models for each sex were run separately. n.s. = not significant.

The third slope pattern included relatively simple slopes that differed between increases and decreases. A linear decrease in iron levels was only seen in female participants’ PuL. An overall increasing log slope across the lifespan was found in female participants’ PuA and LGN and in male participants’ IL, VL, and VPL. Of the thalamic nuclei, female participants’ LP, MGN, PuM, VL, and VPL were not significantly associated with age.

## 4. Discussion

The current cross-sectional study provides strong evidence of an overall non-linear pattern of brain iron across many subcortical regions from adolescence to late adulthood, aligning with previous quantitative MRI studies (Persson et al., 2015; Treit et al., 2021). In detail, we report**e**d that rR2* signal from many of the subcortical ROIs showed nonlinear associations with age, which were the best fit for both female and male participants. Specifically, we revealed that the most prominent trajectory for iron over the lifespan included S-curves.

To our knowledge, we are the first to present lifespan trajectories of brain iron concentrations using fMRI data. Similar previous studies have assessed regional lifespan iron through postmortem histological and chemical analyses (Hallgren & Sourander, 1958, ages: 1-100, *N* = 81), quantitative MRI approaches (Li et al., 2013, ages: 1-83, *N* = 191 and Zhang et al., 2018a, 2018b, ages: 1-83, *N* = 166; Treit et al., 2021, ages: 5-90, *N* = 498). While these studies differed across methodologies, sample sizes, and ROIs they assessed, some similarities were observed. The current results for the trajectories of the pallidum, putamen, and thalamus match those reported in one of the first post-mortem studies by Hallgren and Sourander (1958). Additionally, susceptibility findings from Li et al. (2013) align with those of the current pallidum, putamen, RN, SNpc, and SNpr; while Zhang et al.’s (2018a,b) QSM results align with the current pallidum, RN, SNpc, and SNpr. Importantly, we believe our results include the first reports of lifespan iron trajectories for two subcortical regions, the hippocampus, NA, and VTA, as well as two subregions of the SN.

During early life (∼ <early 20s), iron is in high demand for multiple aspects of neurophysiological development, such as neurotransmitter synthesis (Hidalgo & Núñez, 2007; Kulaszyńska et al., 2024; Piñero & Connor, 2000) and myelination (McCann et al., 2020; Piñero & Connor, 2000; Todorich et al., 2009). The current results show increases in iron for both sexes’ 10 main ROIs, except for female participants’ hippocampus. These findings generally align with those from prior cross-sectional (Parr et al., 2022; Peterson et al., 2018) and longitudinal (Peterson et al., 2018) studies in the developing NA, pallidum, putamen, RN, and overall SN. Lastly, the current caudate trajectory for each sex partially aligns with that of Treit et al. (2021), with initial increasing slopes (F: ∼ <30 years, M: ∼ <50 years).

Throughout middle and late adulthood (> ∼30 years), three overall patterns appeared for the main ROIs, where iron levels either notably increased (putamen and NA), gradually increased (pallidum, RN, SNpc, SNpr, and VTA), or were more complex (caudate, thalamus, and hippocampus). Of the two regions in the first trajectory pattern, the putamen was in agreement with previous reports (Persson et al., 2015), showing a gradual increase from 20 years and older. For the second trajectory, results in pallidum, RN, SNpc, and SNpr iron are aligned with results from Acosta-Cabronero et al. (2016), who found a positive association of susceptibility with age (20-79 years). For the third trajectory pattern, current trajectories found for the thalamus and caudate partially align with results from previous lifespan studies (Hallgren & Sourander, 1958; Treit et al., 2021). For the remaining ROIs, we present novel findings as literature of their normative iron level trajectories across adulthood are presently lacking. The slope patterns for the NA and VTA were simpler (described above); however, the hippocampus was only significant in male individuals, who showed an increase through mid-adulthood (∼ <50 years) followed by a decrease.

The thalamus comprises many nuclei that differ in their functional roles and structural and functional connections (Fama & Sullivan, 2015; Funk et al., 2023). Despite this, there is less literature on differences in iron content between the thalamic nuclei across the healthy lifespan. While it has been suggested that the level of iron is heterogeneous across these nuclei (Treit et al., 2021), our findings do not fully support this. Notably, of the 14 thalamic nuclei used in this study, male participants showed cubic trends similar to their overall thalamus curve for eight, while female participants showed similar slopes for four. Such drastic differences indicate sex-specific rather than nuclei-specific associations in iron-level changes across age that may contribute to prior mixed findings for the overall thalamus.

While this study benefitted from a large sample size, the data used was cross-sectional in nature. With this, we were unable to assess within-individual change over time in our analyses, which has been shown to have significant differences in estimates of age-related neural changes (Di Biase et al., 2023). Therefore, future large, longitudinal datasets will be necessary to assess changes in iron content across the lifespan, particularly in the very early (under 8 years old) and late-life (i.e., above 85) ranges that remain under-researched. However, promising new large and longitudinal studies such as the HEALthy Brain and Child Development (HBCD) Study (Volkow et al., 2024) will likely allow to investigate these trajectories in very early life. Second, as with other in vivo MRI methods currently available to quantify brain iron concentrations, fMRI-based metrics remain indirect, and all results should be interpreted with caution. However, our supplementary analyses demonstrated a strong group-level correspondence between regional averages of the fMRI-derived iron-sensitive signal and both post-mortem data and quantitative in vivo iron metrics. These group-level, regional-average findings suggest that the fMRI-derived iron-sensitive signal (a relative iron quantification computed from fMRI data) is sensitive to and consistent with tissue iron content, rather than providing an individual-level, validated measure of it. An additional limitation concerns the relatively low spatial resolution of the fMRI data (voxel sizes of 3 × 3 × 4.4 mm in the CamCAN sample and 2 × 2 × 2 mm in the local sample), which makes small subcortical structures susceptible to partial-volume contamination from adjacent tissue. As a result, the fMRI-derived iron-sensitive signal extracted from these regions may be influenced by neighboring structures and cerebrospinal fluid. This concern is most pronounced for the smallest ROIs, including the VTA, red nucleus, the substantia nigra subdivisions (SNpc and SNpr), and the individual thalamic nuclei. Accordingly, the results for these regions should be interpreted with additional caution.

We believe the current study includes the largest sample to date to assess the fMRI-derived iron-sensitive signal (relative iron quantification), in subcortical regions. Our results add to the existing literature on subcortical iron content across the lifespan. The findings within sex and across regions suggest that future research should consider more specific ROIs, sex-specific analyses, and additional nonlinear associations when investigating iron levels. The alignment of the current results to those of previous literature provides support for the usage of fMRI to derive a signal that is sensitive to, and consistent with, iron content. While further studies are still necessary to replicate the findings and extend the age range, our study offers an opportunity for more widespread research on brain iron content using a widely applicable fMRI sequence, especially in vulnerable populations who may not be able to tolerate long MRI scans. Lastly, results from the current study demonstrate the need to consider multiple nonlinear associations in age-related changes in iron content across subcortical regions.

## Acknowledgements

This work was supported by the National Institutes of Health (P20 GM144641).

## Data Availability

The first dataset included data from the publicly available the Cambridge Centre for Aging and Neuroscience study (CamCAN). This study used an epidemiologically informed recruitment framework and is available upon request (https://www.cam-can.org/; Shafto et al. 2014; Taylor et al. 2017).

Regional brain iron levels for participants will be made available upon acceptance (a file will be provided at that time).

## Code Availability

The in-home codes used for analyses are available from the corresponding author upon reasonable request.

## Author Contributions

ATM aided in the formulation of overarching research goals, created and analyzed statistical models, wrote the main manuscript text, and prepared tables and figures for main text. DJP: conducted all preprocessing of fMRI data and iron quantification for both datasets. LF assisted in data collection and iron quantification methods. KM conducted experiments and collected data for the local dataset used. CG aided in data management and curation. JJO and TW advised on and verified the proper conductance of statistical methodologies and their interpretations and contributed to manuscript editing. GED assisted in the formulation of overarching research goals, designed study protocol and supervised collection of local dataset, provided study materials, instrumentation, and computing resources, led data management and curation, provided major contributions to manuscript writing and editing, and acquired financial support for the study. All authors reviewed and approved the final manuscript.

## References

Acosta-Cabronero, J., Betts, M. J., Cardenas-Blanco, A., Yang, S., & Nestor, P. J. (2016). In Vivo MRI Mapping of Brain Iron Deposition across the Adult Lifespan. The Journal of Neuroscience, 36(2), 364–374. 10.1523/JNEUROSCI.1907-15.2016

Acosta-Cabronero, J., Williams, G. B., Cardenas-Blanco, A., Arnold, R. J., Lupson, V., & Nestor, P. J. (2013). In Vivo Quantitative Susceptibility Mapping (QSM) in Alzheimer’s Disease. PLoS ONE, 8(11), e81093. 10.1371/journal.pone.0081093

Adisetiyo, V., Gray, K. M., Jensen, J. H., & Helpern, J. A. (2019). Brain iron levels in attention-deficit/hyperactivity disorder normalize as a function of psychostimulant treatment duration. NeuroImage: Clinical, 24, 101993. 10.1016/j.nicl.2019.101993

Ahmadi, S.-A., Bötzel, K., Levin, J., Maiostre, J., Klein, T., Wein, W., Rozanski, V., Dietrich, O., Ertl-Wagner, B., Navab, N., & Plate, A. (2020). Analyzing the co-localization of substantia nigra hyper-echogenicities and iron accumulation in Parkinson’s disease: A multi-modal atlas study with transcranial ultrasound and MRI. NeuroImage: Clinical, 26, 102185. 10.1016/j.nicl.2020.102185

Andersen, H. H., Johnsen, K. B., & Moos, T. (2014). Iron deposits in the chronically inflamed central nervous system and contributes to neurodegeneration. Cellular and Molecular Life Sciences, 71(9), 1607–1622. 10.1007/s00018-013-1509-8

Aquino, D., Bizzi, A., Grisoli, M., Garavaglia, B., Bruzzone, M. G., Nardocci, N., Savoiardo, M., & Chiapparini, L. (2009). Age-related iron deposition in the basal ganglia: Quantitative analysis in healthy subjects. Radiology, 252(1), 165–172. 10.1148/radiol.2522081399

Benjamini, Y., & Hochberg, Y. (1995). Controlling the False Discovery Rate: A Practical and Powerful Approach to Multiple Testing. Journal of the Royal Statistical Society: Series B (Methodological), 57(1), 289–300. 10.1111/j.2517-6161.1995.tb02031.x

Bilgic, B., Pfefferbaum, A., Rohlfing, T., Sullivan, E. V., & Adalsteinsson, E. (2012). MRI Estimates of Brain Iron Concentration in Normal Aging Using Quantitative Susceptibility Mapping. Neuroimage, 59(3), 2625–2635. 10.1016/j.neuroimage.2011.08.077

Brett, M., Markiewicz, C. J., Hanke, M., Côté, M.-A., Cipollini, B., Papadopoulos Orfanos, D., McCarthy, P., Jarecka, D., Cheng, C. P., Larson, E., Halchenko, Y. O., Cottaar, M., Ghosh, S., Wassermann, D., Gerhard, S., Lee, G. R., Baratz, Z., Moloney, B., Wang, H.-T., … freec84. (2025). *nipy/nibabel: 5.3.2* [Computer software]. Zenodo. 10.5281/zenodo.17833216

Burgetova, R., Dusek, P., Burgetova, A., Pudlac, A., Vaneckova, M., Horakova, D., Krasensky, J., Varga, Z., & Lambert, L. (2021). Age-related magnetic susceptibility changes in deep grey matter and cerebral cortex of normal young and middle-aged adults depicted by whole brain analysis. Quantitative Imaging in Medicine and Surgery, 11(9), 3906919–3903919. 10.21037/qims-21-87

Cascone, A. D., Calabro, F., Foran, W., Larsen, B., Nugiel, T., Parr, A. C., Tervo-Clemmens, B., Luna, B., & Cohen, J. R. (2023). Brain tissue iron neurophysiology and its relationship with the cognitive effects of dopaminergic modulation in children with and without ADHD. Developmental Cognitive Neuroscience, 63, 101274. 10.1016/j.dcn.2023.101274

Cortese, S., Azoulay, R., Castellanos, F. X., Chalard, F., Lecendreux, M., Chechin, D., Delorme, R., Sebag, G., Sbarbati, A., Mouren, M.-C., Bernardina, B. D., & Konofal, E. (2012). Brain iron levels in attention-deficit/hyperactivity disorder: A pilot MRI study. The World Journal of Biological Psychiatry: The Official Journal of the World Federation of Societies of Biological Psychiatry, 13(3), 223–231. 10.3109/15622975.2011.570376

Daugherty, A., & Raz, N. (2013). Age-Related Differences in Iron Content of Subcortical Nuclei Observed in vivo: A Meta-Analysis. NeuroImage, 70, 113–121. 10.1016/j.neuroimage.2012.12.040

Di Martino, A., O’Connor, D., Chen, B., Alaerts, K., Anderson, J. S., Assaf, M., Balsters, J. H., Baxter, L., Beggiato, A., Bernaerts, S., Blanken, L. M. E., Bookheimer, S. Y., Braden, B. B., Byrge, L., Castellanos, F. X., Dapretto, M., Delorme, R., Fair, D. A., Fishman, I., … Milham, M. P. (2017). Enhancing studies of the connectome in autism using the autism brain imaging data exchange II. Scientific Data, 4, 170010. 10.1038/sdata.2017.10

Di Martino, A., Yan, C.-G., Li, Q., Denio, E., Castellanos, F. X., Alaerts, K., Anderson, J. S., Assaf, M., Bookheimer, S. Y., Dapretto, M., Deen, B., Delmonte, S., Dinstein, I., Ertl-Wagner, B., Fair, D. A., Gallagher, L., Kennedy, D. P., Keown, C. L., Keysers, C., … Milham, M. P. (2014). The Autism Brain Imaging Data Exchange: Towards Large-Scale Evaluation of the Intrinsic Brain Architecture in Autism. Molecular Psychiatry, 19(6), 659–667. 10.1038/mp.2013.78

Di Biase, M. A., Tian, Y. E., Bethlehem, R. A. I., Seidlitz, J., Alexander-Bloch, Aaron. F., Yeo, B. T. T., & Zalesky, A. (2023). Mapping human brain charts cross-sectionally and longitudinally. Proceedings of the National Academy of Sciences, 120(20), e2216798120. 10.1073/pnas.2216798120

Dixon, S. J., & Stockwell, B. R. (2014). The role of iron and reactive oxygen species in cell death. Nature Chemical Biology, 10(1), 9–17. 10.1038/nchembio.1416

Doucet, G. E., Kruse, J. A., Mertens, A., Goldsmith, C., Eden, N. M., Oleson, J., & McGregor, K. K. (2025). Subcortical brain iron and its link to verbal memory in children with developmental language disorder. Brain and Language, 261, 105531. 10.1016/j.bandl.2024.105531

Elam, J. S., Glasser, M. F., Harms, M. P., Sotiropoulos, S. N., Andersson, J. L. R., Burgess, G. C., Curtiss, S. W., Oostenveld, R., Larson-Prior, L. J., Schoffelen, J.-M., Hodge, M. R., Cler, E. A., Marcus, D. M., Barch, D. M., Yacoub, E., Smith, S. M., Ugurbil, K., & Van Essen, D. C. (2021). The Human Connectome Project: A retrospective. NeuroImage, 244, 118543. 10.1016/j.neuroimage.2021.118543

Fama, R., & Sullivan, E. V. (2015). Thalamic structures and associated cognitive functions: Relations with age and aging. Neuroscience and Biobehavioral Reviews, 54, 29–37. 10.1016/j.neubiorev.2015.03.008

Feng, X., Deistung, A., & Reichenbach, J. R. (2018). Quantitative susceptibility mapping (QSM) and R2* in the human brain at 3T: Evaluation of intra-scanner repeatability. Zeitschrift Fur Medizinische Physik, 28(1), 36–48. 10.1016/j.zemedi.2017.05.003

Flannery, J. S., Parr, A. C., Lindquist, K. A., & Telzer, E. H. (2025). Developmental changes in dopamine-related neurophysiology and associations with adolescent substance use and incentive-boosted cognitive control. Developmental Cognitive Neuroscience, 75, 101594. 10.1016/j.dcn.2025.101594

Fortin, J.-P., Cullen, N., Sheline, Y. I., Taylor, W. D., Aselcioglu, I., Cook, P. A., Adams, P., Cooper, C., Fava, M., McGrath, P. J., McInnis, M., Phillips, M. L., Trivedi, M. H., Weissman, M. M., & Shinohara, R. T. (2018). Harmonization of cortical thickness measurements across scanners and sites. NeuroImage, 167, 104–120. 10.1016/j.neuroimage.2017.11.024

Fortin, J.-P., Parker, D., Tunç, B., Watanabe, T., Elliott, M. A., Ruparel, K., Roalf, D. R., Satterthwaite, T. D., Gur, R. C., Gur, R. E., Schultz, R. T., Verma, R., & Shinohara, R. T. (2017). Harmonization of multi-site diffusion tensor imaging data. NeuroImage, 161, 149–170. 10.1016/j.neuroimage.2017.08.047

Funk, A. T., Hassan, A. A. O., Brüggemann, N., Sharma, N., Breiter, H. C., Blood, A. J., & Waugh, J. L. (2023). In humans, striato-pallido-thalamic projections are largely segregated by their origin in either the striosome-like or matrix-like compartments. Frontiers in Neuroscience, 17, 1178473. 10.3389/fnins.2023.1178473

Ghadery, C., Pirpamer, L., Hofer, E., Langkammer, C., Petrovic, K., Loitfelder, M., Schwingenschuh, P., Seiler, S., Duering, M., Jouvent, E., Schmidt, H., Fazekas, F., Mangin, J.-F., Chabriat, H., Dichgans, M., Ropele, S., & Schmidt, R. (2015). R2* mapping for brain iron: Associations with cognition in normal aging. Neurobiology of Aging, 36(2), 925–932. 10.1016/j.neurobiolaging.2014.09.013

Glasser, M. F., Smith, S. M., Marcus, D. S., Andersson, J. L. R., Auerbach, E. J., Behrens, T. E. J., Coalson, T. S., Harms, M. P., Jenkinson, M., Moeller, S., Robinson, E. C., Sotiropoulos, S. N., Xu, J., Yacoub, E., Ugurbil, K., & Van Essen, D. C. (2016). The Human Connectome Project’s neuroimaging approach. Nature Neuroscience, 19(9), 1175–1187. 10.1038/nn.4361

Glasser, M. F., Sotiropoulos, S. N., Wilson, J. A., Coalson, T. S., Fischl, B., Andersson, J. L., Xu, J., Jbabdi, S., Webster, M., Polimeni, J. R., Van Essen, D. C., Jenkinson, M., & WU-Minn HCP Consortium. (2013). The minimal preprocessing pipelines for the Human Connectome Project. NeuroImage, 80, 105–124. 10.1016/j.neuroimage.2013.04.127

Haacke, E. M., Miao, Y., Liu, M., Habib, C. A., Katkuri, Y., Liu, T., Yang, Z., Lang, Z., Hu, J., & Wu, J. (2010). Correlation of change in R2* and phase with putative iron content in deep gray matter of healthy adults. Journal of Magnetic Resonance Imaging : JMRI, 32(3), 561–576. 10.1002/jmri.22293

Hallgren, B., & Sourander, P. (1958). The Effect of Age on the Non-Haemin Iron in the Human Brain. Journal of Neurochemistry, 3(1), 41–51. 10.1111/j.1471-4159.1958.tb12607.x

Harris, C. R., Millman, K. J., van der Walt, S. J., Gommers, R., Virtanen, P., Cournapeau, D., Wieser, E., Taylor, J., Berg, S., Smith, N. J., Kern, R., Picus, M., Hoyer, S., van Kerkwijk, M. H., Brett, M., Haldane, A., del Río, J. F., Wiebe, M., Peterson, P., … Oliphant, T. E. (2020). Array programming with NumPy. Nature, 585(7825), 357–362. 10.1038/s41586-020-2649-2

Hasaneen, B. M., Sarhan, M., Samir, S., ELAssmy, M., Sakrana, A. A., & Ashamalla, G. A. (2017). T2* magnetic resonance imaging: A non-invasive biomarker of brain iron content in children with attention-deficit/hyperactivity disorder. The Egyptian Journal of Radiology and Nuclear Medicine, 48(1), 161–167. 10.1016/j.ejrnm.2016.08.001

Hect, J. L., Daugherty, A. M., Hermez, K. M., & Thomason, M. E. (2018). Developmental variation in regional brain iron and its relation to cognitive functions in childhood. Developmental Cognitive Neuroscience, 34, 18–26. 10.1016/j.dcn.2018.05.004

Hidalgo, C., & Núñez, M. T. (2007). Calcium, iron and neuronal function. IUBMB Life, 59(4–5), 280–285. 10.1080/15216540701222906

Kagerer, S. M., Vionnet, L., van Bergen, J. M. G., Meyer, R., Gietl, A. F., Pruessmann, K. P., Hock, C., & Unschuld, P. G. (2025). Hippocampal iron patterns in aging and mild cognitive impairment. Frontiers in Aging Neuroscience, 17, 1598859. 10.3389/fnagi.2025.1598859

Kulaszynska, M., Kwiatkowski, S., & Skonieczna-Żydecka, K. (2024). The Iron Metabolism with a Specific Focus on the Functioning of the Nervous System. Biomedicines, 12(3), 595. 10.3390/biomedicines12030595

Langkammer, C., Krebs, N., Goessler, W., Scheurer, E., Ebner, F., Yen, K., Fazekas, F., & Ropele, S. (2010). Quantitative MR imaging of brain iron: A postmortem validation study. Radiology, 257(2), 455–462. 10.1148/radiol.10100495

Langkammer, C., Schweser, F., Krebs, N., Deistung, A., Goessler, W., Scheurer, E., Sommer, K., Reishofer, G., Yen, K., Fazekas, F., Ropele, S., & Reichenbach, J. R. (2012). Quantitative susceptibility mapping (QSM) as a means to measure brain iron? A post mortem validation study. Neuroimage, 62(3–2), 1593–1599. 10.1016/j.neuroimage.2012.05.049

Larsen, B., Bourque, J., Moore, T. M., Adebimpe, A., Calkins, M. E., Elliott, M. A., Gur, R. C., Gur, R. E., Moberg, P. J., Roalf, D. R., Ruparel, K., Turetsky, B. I., Vandekar, S. N., Wolf, D. H., Shinohara, R. T., & Satterthwaite, T. D. (2020). Longitudinal Development of Brain Iron Is Linked to Cognition in Youth. The Journal of Neuroscience, 40(9), 1810–1818. 10.1523/JNEUROSCI.2434-19.2020

Larsen, B., & Luna, B. (2015). In vivo evidence of neurophysiological maturation of the human adolescent striatum. Developmental Cognitive Neuroscience, 12, 74–85. 10.1016/j.dcn.2014.12.003

Li, G., Tong, R., Zhang, M., Gillen, K. M., Jiang, W., Du, Y., Wang, Y., & Li, J. (2023). Age-dependent changes in brain iron deposition and volume in deep gray matter nuclei using quantitative susceptibility mapping. NeuroImage, 269, 119923. 10.1016/j.neuroimage.2023.119923

Li, W., Wu, B., Batrachenko, A., Bancroft-Wu, V, Morey, R. A., Shashi, V., Langkammer, C., De Bellis, M. D., Ropele, S., Song, A. W., & Liu, C. (2013). Differential developmental trajectories of magnetic susceptibility in human brain gray and white matter over the lifespan. Human Brain Mapping, 35(6), 2698–2713. 10.1002/hbm.22360

López-Aguirre, M., Balzano, T., Monje, M. H. G., Esteban-García, N., Martínez-Fernández, R., Del Rey, N. L., Ciorraga, M., Sánchez-Ferro, A., Trigo-Damas, I., Blesa, J., Obeso, J. A., & Pineda-Pardo, J. A. (2025). Nigrostriatal iron accumulation in the progression of Parkinson’s disease. Npj Parkinson’s Disease, 11(1), 72. 10.1038/s41531-025-00911-6

Madden, D. J., & Merenstein, J. L. (2023). Quantitative susceptibility mapping of brain iron in healthy aging and cognition. NeuroImage, 282, 120401. 10.1016/j.neuroimage.2023.120401

Marek, K., Jennings, D., Lasch, S., Siderowf, A., Tanner, C., Simuni, T., Coffey, C., Kieburtz, K., Flagg, E., Chowdhury, S., Poewe, W., Mollenhauer, B., Klinik, P.-E., Sherer, T., Frasier, M., Meunier, C., Rudolph, A., Casaceli, C., Seibyl, J., … Taylor, P. (2011). The Parkinson Progression Marker Initiative (PPMI). Progress in Neurobiology, 95(4), 629–635. 10.1016/j.pneurobio.2011.09.005

McCann, S., Perapoch Amadó, M., & Moore, S. E. (2020). The Role of Iron in Brain Development: A Systematic Review. Nutrients, 12(7), 2001. 10.3390/nu12072001

Mieling, M., Wiskow, C., & Bunzeck, N. (2025). Globus Pallidus Iron Relates to Cognitive Impairment in Alzheimer’s Disease: Evidence From MRI-Based Meta-Analysis. Annals of the New York Academy of Sciences. 10.1111/nyas.70078

Parr, A. C., Calabro, F., Tervo-Clemmens, B., Larsen, B., Foran, W., & Luna, B. (2022). Contributions of dopamine-related basal ganglia neurophysiology to the developmental effects of incentives on inhibitory control. Developmental Cognitive Neuroscience, 54, 101100. 10.1016/j.dcn.2022.101100

Parr, A. C., Ojha, A., Petrie, D. J., Calabro, F. J., Tervo-Clemmens, B., Foran, W., Fitzgerald, D., Tapert, S. F., Nooner, K., Thompson, W., Goldston, D. B., Clark, D., & Luna, B. (2026). Developmental variation in basal ganglia tissue iron, neurocognitive functioning, and impulsivity is associated with substance use trajectories in youth. Nature Communications, 17(1), 4861. 10.1038/s41467-026-73611-1

Péran, P., Cherubini, A., Luccichenti, G., Hagberg, G., Démonet, J., Rascol, O., Celsis, P., Caltagirone, C., Spalletta, G., & Sabatini, U. (2009). Volume and iron content in basal ganglia and thalamus. Human Brain Mapping, 30(8), 2667–2675. 10.1002/hbm.20698

Persson, N., Wu, J., Zhang, Q., Liu, T., Shen, J., Bao, R., Ni, M., Liu, T., Wang, Y., & Spincemaille, P. (2015). Age and Sex Related Differences in Subcortical Brain Iron Concentrations among Healthy Adults. NeuroImage, 122, 385–398. 10.1016/j.neuroimage.2015.07.050

Peterson, E. T., Kwon, D., Luna, B., Larsen, B., Prouty, D., De Bellis, M. D., Voyvodic, J., Liu, C., Li, W., Pohl, K. M., Sullivan, E. V., & Pfefferbaum, A. (2018). Distribution of brain iron accrual in adolescence: Evidence from cross-sectional and longitudinal analysis. Human Brain Mapping, 40(5), 1480–1495. 10.1002/hbm.24461

Petok, J. R., Merenstein, J. L., & Bennett, I. J. (2024). Iron content affects age group differences in associative learning-related fMRI activity. NeuroImage, 285, 120478. 10.1016/j.neuroimage.2023.120478

Pfefferbaum, A., Adalsteinsson, E., Rohlfing, T., & Sullivan, E. V. (2009). MRI estimates of brain iron concentration in normal aging: Comparison of field-dependent (FDRI) and phase (SWI) methods. NeuroImage, 47(2), 493–500. 10.1016/j.neuroimage.2009.05.006

Piñero, D. J., & Connor, J. R. (2000). Iron in the Brain: An Important Contributor in Normal and Diseased States. The Neuroscientist, 6(6), 435–453. 10.1177/107385840000600607

Price, R. B., Tervo-Clemmens, B. C., Panny, B., Degutis, M., Griffo, A., & Woody, M. (2021). Biobehavioral correlates of an fMRI index of striatal tissue iron in depressed patients. Translational Psychiatry, 11(1), 448. 10.1038/s41398-021-01553-x

Ramos, P., Santos, A., Pinto, N. R., Mendes, R., Magalhães, T., & Almeida, A. (2014). Iron levels in the human brain: A post-mortem study of anatomical region differences and age-related changes. Journal of Trace Elements in Medicine and Biology, 28(1), 13–17. 10.1016/j.jtemb.2013.08.001

Rolls, E. T., Huang, C.-C., Lin, C.-P., Feng, J., & Joliot, M. (2020). Automated anatomical labelling atlas 3. NeuroImage, 206, 116189. 10.1016/j.neuroimage.2019.116189

Schulze, M., Coghill, D., Lux, S., Philipsen, A., & Silk, T. (2025). Assessing Brain Iron and Its Relationship to Cognition and Comorbidity in Children With Attention-Deficit/Hyperactivity Disorder With Quantitative Susceptibility Mapping. Biological Psychiatry: Cognitive Neuroscience and Neuroimaging, 10(6), 597–606. 10.1016/j.bpsc.2024.08.015

Shafto, M. A., Tyler, L. K., Dixon, M., Taylor, J. R., Rowe, J. B., Cusack, R., Calder, A. J., Marslen-Wilson, W. D., Duncan, J., Dalgleish, T., Henson, R. N., Brayne, C., Matthews, F. E., & Cam-CAN. (2014). The Cambridge Centre for Ageing and Neuroscience (Cam-CAN) study protocol: A cross-sectional, lifespan, multidisciplinary examination of healthy cognitive ageing. BMC Neurology, 14(1), 204. 10.1186/s12883-014-0204-1

Sonnenschein, S. F., Parr, A. C., Larsen, B., Calabro, F. J., Foran, W., Eack, S. M., Luna, B., & Sarpal, D. K. (2022). Subcortical brain iron deposition in individuals with schizophrenia. Journal of Psychiatric Research, 151, 272–278. 10.1016/j.jpsychires.2022.04.013

Spence, H., McNeil, C. J., & Waiter, G. D. (2020). The impact of brain iron accumulation on cognition: A systematic review. PLoS ONE, 15(10), e0240697. 10.1371/journal.pone.0240697

Spincemaille, P., Liu, Z., Zhang, S., Kovanlikaya, I., Ippoliti, M., Makowski, M., Watts, R., de Rochefort, L., Venkatraman, V., Desmond, P., Santin, M. D., Lehéricy, S., Kopell, B. H., Péran, P., & Wang, Y. (2019). Clinical integration of automated processing for brain quantitative susceptibility mapping: Multi-site reproducibility and single-site robustness. Journal of Neuroimaging : Official Journal of the American Society of Neuroimaging, 29(6), 689–698. 10.1111/jon.12658

Spotorno, N., Acosta-Cabronero, J., Stomrud, E., Lampinen, B., Strandberg, O. T., Van Westen, D., & Hansson, O. (2020). Relationship between cortical iron and tau aggregation in Alzheimer’s disease. Brain, 143(5), 1341–1349. 10.1093/brain/awaa089

Taylor, J. R., Williams, N., Cusack, R., Auer, T., Shafto, M. A., Dixon, M., Tyler, L. K., Cam-CAN, & Henson, R. N. (2017). The Cambridge Centre for Ageing and Neuroscience (Cam-CAN) data repository: Structural and functional MRI, MEG, and cognitive data from a cross-sectional adult lifespan sample. Neuroimage, 144(Pt B), 262–269. 10.1016/j.neuroimage.2015.09.018

Todorich, B., Pasquini, J. M., Garcia, C. I., Paez, P. M., & Connor, J. R. (2009). Oligodendrocytes and myelination: The role of iron. Glia, 57(5), 467–478. 10.1002/glia.20784

Treit, S., Naji, N., Seres, P., Rickard, J., Stolz, E., Wilman, A. H., & Beaulieu, C. (2021). R2* and quantitative susceptibility mapping in deep gray matter of 498 healthy controls from 5 to 90 years. Human Brain Mapping, 42(14), 4597–4610. 10.1002/hbm.25569

Volkow, N. D., Gordon, J. A., Bianchi, D. W., Chiang, M. F., Clayton, J. A., Klein, W. M., Koob, G. F., Koroshetz, W. J., Pérez-Stable, E. J., Simoni, J. M., Tromberg, B. J., Woychik, R. P., Hommer, R., Spotts, E. L., Xu, B., Zehr, J. L., Cole, K. M., Dowling, G. J., Freund, M. P., … Weiss, S. R. B. (2024). The HEALthy Brain and Child Development Study (HBCD): NIH collaboration to understand the impacts of prenatal and early life experiences on brain development. Developmental Cognitive Neuroscience, 69, 101423. 10.1016/j.dcn.2024.101423

Wang, Z.-L., Yuan, L., Li, W., & Li, J.-Y. (2022). Ferroptosis in Parkinson’s disease: Glia-neuron crosstalk. Trends in Molecular Medicine, 28(4), 258–269. 10.1016/j.molmed.2022.02.003

Xu, X., Wang, Q., & Zhang, M. (2008). Age, gender, and hemispheric differences in iron deposition in the human brain: An in vivo MRI study. NeuroImage, 40(1), 35–42. 10.1016/j.neuroimage.2007.11.017

Yan, S.-Q., Sun, J.-Z., Yan, Y.-Q., Wang, H., & Lou, M. (2012). Evaluation of Brain Iron Content Based on Magnetic Resonance Imaging (MRI): Comparison among Phase Value, R2* and Magnitude Signal Intensity. PLoS ONE, 7(2), e31748. 10.1371/journal.pone.0031748

Zeineh, M. M., Chen, Y., Kitzler, H. H., Hammond, R., Vogel, H., & Rutt, B. K. (2015). Activated iron-containing microglia in the human hippocampus identified by magnetic resonance imaging in Alzheimer disease. Neurobiology of Aging, 36(9), 2483–2500. 10.1016/j.neurobiolaging.2015.05.022

Zhang, Y., Wei, H., Cronin, M. J., He, N., Yan, F., & Liu, C. (2018a). Longitudinal Atlas for Normative Human Brain Development and Aging over the Lifespan using Quantitative Susceptibility Mapping. NeuroImage, 171, 176–189. 10.1016/j.neuroimage.2018.01.008

Zhang, Y., Wei, H., Cronin, M. J., He, N., Yan, F., & Liu, C. (2018b). Longitudinal data for magnetic susceptibility of normative human brain development and aging over the lifespan. Data in Brief, 20, 623–631. 10.1016/j.dib.2018.06.005

