## Supplementary Materials for "Patterns of subcortical tissue iron levels across the lifespan measured with functional MRI"

**Tables**

Supplementary Table S1: All Tested Regression Models for Main 10 ROIs

| ROI | Slope | Sex | Variable | *F* | df1 | df2 | *p_FDR_* | *R^2^* | *b* | *SE* | *t* | *p_FDR_* |
| --- | --- | --- | --- | --- | --- | --- | --- | --- | --- | --- | --- | --- |
| Caudate | Linear | F | Model | 32.91 | 1 | 443 | 6.84x10^-8^ | 0.07 |  | | | |
|  |  |  | Age |  | | | | | -4.00x10^-4^ | 6.37x10^-5^ | -5.74 | 6.84x10^-8^ |
|  |  | M | Model | 3.05 | 1 | 419 | .102 | 0.01 |  | | | |
|  |  |  | Age |  | | | | | -1.00x10^-4^ | 6.77x10^-5^ | -1.75 | .100 |
|  | Quadratic | F | Model | 29.25 | 2 | 442 | 5.90x10^-12^ | 0.12 |  | | | |
|  |  |  | Age |  | | | | | -0.01 | 0.001 | -5.38 | 4.29x10^-7^ |
|  |  |  | Age^2^ |  | | | | | -0.01 | 0.002 | -4.89 | 4.77x10^-6^ |
|  |  | M | Model | 7.72 | 2 | 418 | 8.41x10^-4^ | 0.04 |  | | | |
|  |  |  | Age |  | | | | | -0.003 | 0.002 | -1.84 | .086 |
|  |  |  | Age^2^ |  | | | | | -0.01 | 0.002 | -3.26 | .002 |
|  | Cubic | F | Model | 24.10 | 3 | 441 | 1.09x10^-13^ | 0.14 |  | | | |
|  |  |  | Age |  | | | | | -0.02 | 0.003 | -5.41 | 3.83x10^-7^ |
|  |  |  | Age^2^ |  | | | | | -0.01 | 0.002 | -5.70 | 8.06x10^-8^ |
|  |  |  | Age^3^ |  | | | | | 0.01 | 0.002 | 3.51 | 8.95x10^-4^ |
|  |  | M | Model | 6.50 | 3 | 417 | 4.52x10^-4^ | 0.05 |  | | | |
|  |  |  | Age |  | | | | | -0.01 | 0.004 | -2.57 | .015 |
|  |  |  | Age^2^ |  | | | | | -0.01 | 0.002 | -3.66 | 4.80x10^-4^ |
|  |  |  | Age^3^ |  | | | | | 0.004 | 0.002 | 2.00 | .061 |
|  | Log | F | Model | 18.30 | 1 | 443 | 6.26x10^-5^ | 0.04 |  | | | |
|  |  |  | Age |  | | | | | -0.01 | 0.003 | -4.28 | 6.26x10^-5^ |
|  |  | M | Model | 0.55 | 1 | 419 | .514 | 0.001 |  | | | |
|  |  |  | Age |  | | | | | -0.002 | 0.003 | -0.74 | .514 |
|  | S-curve | F | Model | 6.36 | 1 | 443 | .018 | 0.01 |  | | | |
|  |  |  | Age |  | | | | | 0.26 | 0.10 | 2.52 | .018 |
|  |  | M | Model | 0.0002 | 1 | 419 | .990 | 4.12x10^-7^ |  | | | |
|  |  |  | Age |  | | | | | -0.001 | 0.09 | -0.01 | .990 |
| Hippocampus | Linear | F | Model | 0.36 | 1 | 433 | .582 | 0.001 |  | | | |
|  |  |  | Age |  | | | | | 6.16x10^-5^ | 0.0001 | 0.60 | .582 |
|  |  | M | Model | 0.06 | 1 | 416 | .839 | 0.0001 |  | | | |
|  |  |  | Age |  | | | | | 2.15x10^-5^ | 8.78x10^-5^ | 0.25 | .839 |
|  | Quadratic | F | Model | 0.85 | 2 | 432 | .483 | 0.004 |  | | | |
|  |  |  | Age |  | | | | | 0.002 | 0.002 | 0.71 | .523 |
|  |  |  | Age^2^ |  | | | | | -0.003 | 0.003 | -1.16 | .299 |
|  |  | M | Model | 3.93 | 2 | 415 | .029 | 0.02 |  | | | |
|  |  |  | Age |  | | | | | 0.001 | 0.002 | 0.54 | .626 |
|  |  |  | Age^2^ |  | | | | | -0.006 | 0.002 | -2.79 | .008 |
|  | Cubic | F | Model | 0.62 | 3 | 431 | .634 | 0.004 |  | | | |
|  |  |  | Age |  | | | | | 0.004 | 0.01 | 0.64 | .567 |
|  |  |  | Age^2^ |  | | | | | -0.003 | 0.003 | -1.01 | .368 |
|  |  |  | Age^3^ |  | | | | | -0.001 | 0.003 | -0.39 | .722 |
|  |  | M | Model | 3.61 | 3 | 414 | .019 | 0.03 |  | | | |
|  |  |  | Age |  | | | | | -0.01 | 0.01 | -1.35 | .211 |
|  |  |  | Age^2^ |  | | | | | -0.01 | 0.002 | -3.14 | .003 |
|  |  |  | Age^3^ |  | | | | | 0.01 | 0.003 | 1.71 | .106 |
|  | Log | F | Model | 0.73 | 1 | 433 | .449 | 0.002 |  | | | |
|  |  |  | Age |  | | | | | 0.004 | 0.004 | 0.85 | .449 |
|  |  | M | Model | 0.95 | 1 | 416 | .379 | 0.002 |  | | | |
|  |  |  | Age |  | | | | | 0.004 | 0.004 | 0.98 | .379 |
|  | S-curve | F | Model | 1.30 | 1 | 433 | .300 | 0.003 |  | | | |
|  |  |  | Age |  | | | | | -0.17 | 0.15 | -1.14 | .300 |
|  |  | M | Model | 3.28 | 1 | 416 | .089 | 0.01 |  | | | |
|  |  |  | Age |  | | | | | -0.24 | 0.13 | -1.81 | .089 |
| NA | Linear | F | Model | 13.01 | 1 | 422 | 6.95x10^-4^ | 0.03 |  | | | |
|  |  |  | Age |  | | | | | 0.001 | 0.0002 | 3.61 | 6.95x10^-4^ |
|  |  | M | Model | 20.82 | 1 | 404 | 1.32x10^-5^ | 0.05 |  | | | |
|  |  |  | Age |  | | | | | 0.001 | 0.0002 | 4.56 | 1.32x10^-5^ |
|  | Quadratic | F | Model | 7.89 | 2 | 421 | 8.00x10^-4^ | 0.04 |  | | | |
|  |  |  | Age |  | | | | | 0.01 | 0.004 | 3.40 | .001 |
|  |  |  | Age^2^ |  | | | | | 0.01 | 0.01 | 1.65 | .125 |
|  |  | M | Model | 10.68 | 2 | 403 | 5.57x10^-5^ | 0.05 |  | | | |
|  |  |  | Age |  | | | | | 0.02 | 0.004 | 4.61 | 1.10x10^-5^ |
|  |  |  | Age^2^ |  | | | | | -0.003 | 0.05 | -0.74 | .514 |
|  | Cubic | F | Model | 6.72 | 3 | 420 | 4.46x10^-4^ | 0.05 |  | | | |
|  |  |  | Age |  | | | | | 0.03 | 0.01 | 3.27 | .002 |
|  |  |  | Age^2^ |  | | | | | 0.01 | 0.004 | 2.18 | .041 |
|  |  |  | Age^3^ |  | | | | | -0.01 | 0.01 | -2.06 | .053 |
|  |  | M | Model | 8.77 | 3 | 402 | 2.30x10^-5^ | 0.06 |  | | | |
|  |  |  | Age |  | | | | | -0.001 | 0.01 | -0.10 | .938 |
|  |  |  | Age^2^ |  | | | | | -0.01 | 0.01 | -1.27 | .242 |
|  |  |  | Age^3^ |  | | | | | 0.01 | 0.01 | 2.18 | .041 |
|  | Log | F | Model | 9.11 | 1 | 422 | .004 | 0.02 |  | | | |
|  |  |  | Age |  | | | | | 0.02 | 0.01 | 3.02 | .004 |
|  |  | M | Model | 23.23 | 1 | 404 | 4.42x10^-6^ | 0.05 |  | | | |
|  |  |  | Age |  | | | | | 0.04 | 0.01 | 4.82 | 4.42x10^-6^ |
|  | S-curve | F | Model | 5.51 | 1 | 422 | .027 | 0.01 |  | | | |
|  |  |  | Age |  | | | | | -0.54 | 0.23 | -2.35 | .027 |
|  |  | M | Model | 28.16 | 1 | 404 | 4.43x10^-7^ | 0.07 |  | | | |
|  |  |  | Age |  | | | | | -1.36 | 0.26 | -5.31 | 4.43x10^-7^ |
| Pallidum | Linear | F | Model | 47.13 | 1 | 429 | 1.05x10^-10^ | 0.10 |  | | | |
|  |  |  | Age |  | | | | | 0.002 | 2x10^-4^ | 6.86 | 1.05x10^-10^ |
|  |  | M | Model | 103.27 | 1 | 412 | 5.01x10^-21^ | 0.20 |  | | | |
|  |  |  | Age |  | | | | | 0.003 | 3x10^-4^ | 10.16 | 5.01x10^-21^ |
|  | Quadratic | F | Model | 36.97 | 2 | 428 | 1.02x10^-14^ | 0.15 |  | | | |
|  |  |  | Age |  | | | | | 0.03 | 0.01 | 6.82 | 1.33x10^-10^ |
|  |  |  | Age^2^ |  | | | | | -0.02 | 0.01 | -3.67 | 5.83x10^-4^ |
|  |  | M | Model | 58.76 | 2 | 411 | 2.74x10^-22^ | 0.22 |  | | | |
|  |  |  | Age |  | | | | | 0.06 | 0.01 | 9.37 | 2.57x10^-18^ |
|  |  |  | Age^2^ |  | | | | | -0.01 | 0.01 | -2.17 | .042 |
|  | Cubic | F | Model | 23.85 | 3 | 427 | 1.54x10^-13^ | 0.14 |  | | | |
|  |  |  | Age |  | | | | | 0.02 | 0.01 | 1.84 | .086 |
|  |  |  | Age^2^ |  | | | | | -0.02 | 0.01 | -3.53 | 8.32x10^-4^ |
|  |  |  | Age^3^ |  | | | | | 0.01 | 0.01 | 0.86 | .449 |
|  |  | M | Model | 44.20 | 3 | 410 | 7.41x10^-24^ | 0.24 |  | | | |
|  |  |  | Age |  | | | | | 0.03 | 0.01 | 2.11 | .047 |
|  |  |  | Age^2^ |  | | | | | -0.01 | 0.01 | -1.81 | .089 |
|  |  |  | Age^3^ |  | | | | | 0.02 | 0.01 | 1.99 | .062 |
|  | Log | F | Model | 68.57 | 1 | 429 | 1.02x10^-14^ | 0.14 |  | | | |
|  |  |  | Age |  | | | | | 0.07 | 0.01 | 8.28 | 1.02x10^-14^ |
|  |  | M | Model | 134.04 | 1 | 412 | 4.31x10^-26^ | 0.25 |  | | | |
|  |  |  | Age |  | | | | | 0.10 | 0.01 | 11.58 | 4.31x10^-26^ |
|  | S-curve | F | Model | 98.86 | 1 | 429 | 9.17x10^-20^ | 0.19 |  | | | |
|  |  |  | Age |  | | | | | -1.73 | 0.17 | -9.94 | 9.17x10^-20^ |
|  |  | M | Model | 161.66 | 1 | 412 | 2.59x10^-30^ | 0.28 |  | | | |
|  |  |  | Age |  | | | | | -2.35 | 0.19 | -12.71 | 2.59x10^-30^ |
| Putamen | Linear | F | Model | 76.68 | 1 | 427 | 4.11x10^-16^ | 0.15 |  | | | |
|  |  |  | Age |  | | | | | 0.001 | 1x10^-4^ | 8.76 | 4.11x10^-16^ |
|  |  | M | Model | 190.87 | 1 | 407 | 4.66x10^-34^ | 0.32 |  | | | |
|  |  |  | Age |  | | | | | 0.002 | 1x10^-4^ | 13.82 | 4.66x10^-34^ |
|  | Quadratic | F | Model | 39.12 | 2 | 426 | 2.06x10^-15^ | 0.16 |  | | | |
|  |  |  | Age |  | | | | | 0.02 | 0.003 | 8.51 | 2.29x10^-15^ |
|  |  |  | Age^2^ |  | | | | | -0.001 | 0.003 | -0.44 | .684 |
|  |  | M | Model | 93.84 | 2 | 406 | 8.24x10^-33^ | 0.32 |  | | | |
|  |  |  | Age |  | | | | | 0.04 | 0.003 | 12.77 | 2.40x10^-30^ |
|  |  |  | Age^2^ |  | | | | | 0.002 | 0.003 | 0.68 | .543 |
|  | Cubic | F | Model | 25.21 | 3 | 425 | 3.10x10^-14^ | 0.15 |  | | | |
|  |  |  | Age |  | | | | | 0.02 | 0.006 | 3.61 | 6.95x10^-4^ |
|  |  |  | Age^2^ |  | | | | | -0.001 | 0.003 | -0.36 | .736 |
|  |  |  | Age^3^ |  | | | | | -0.001 | 0.003 | -0.15 | .888 |
|  |  | M | Model | 62.58 | 3 | 405 | 6.31x10^-32^ | 0.32 |  | | | |
|  |  |  | Age |  | | | | | 0.04 | 0.01 | 5.93 | 1.80x10^-8^ |
|  |  |  | Age^2^ |  | | | | | 0.002 | 0.003 | 0.58 | .607 |
|  |  |  | Age^3^ |  | | | | | -0.001 | 0.004 | -0.21 | .855 |
|  | Log | F | Model | 85.90 | 1 | 427 | 1.02x10^-17^ | 0.17 |  | | | |
|  |  |  | Age |  | | | | | 0.04 | 0.004 | 9.27 | 1.02x10^-17^ |
|  |  | M | Model | 184.21 | 1 | 407 | 2.30x10^-33^ | 0.32 |  | | | |
|  |  |  | Age |  | | | | | 0.06 | 0.004 | 13.57 | 2.30x10^-33^ |
|  | S-curve | F | Model | 89.73 | 1 | 427 | 2.46x10^-18^ | 0.17 |  | | | |
|  |  |  | Age |  | | | | | -1.03 | 0.11 | -9.47 | 2.46x10^-18^ |
|  |  | M | Model | 148.21 | 1 | 407 | 3.23x10^-28^ | 0.27 |  | | | |
|  |  |  | Age |  | | | | | -1.48 | 0.12 | -12.17 | 3.23x10^-28^ |
| RN | Linear | F | Model | 8.05 | 1 | 424 | .008 | 0.02 |  | | | |
|  |  |  | Age |  | | | | | 0.001 | 2x10^-4^ | 2.84 | .008 |
|  |  | M | Model | 38.27 | 1 | 411 | 4.50x10^-9^ | 0.09 |  | | | |
|  |  |  | Age |  | | | | | 0.001 | 2x10^-4^ | 6.19 | 4.50x10^-9^ |
|  | Quadratic | F | Model | 10.50 | 2 | 423 | 9.18x10^-5^ | 0.05 |  | | | |
|  |  |  | Age |  | | | | | 0.01 | 0.004 | 3.23 | .002 |
|  |  |  | Age^2^ |  | | | | | -0.02 | 0.01 | -3.57 | 7.52x10^-4^ |
|  |  | M | Model | 32.52 | 2 | 410 | 2.90x10^-13^ | 0.14 |  | | | |
|  |  |  | Age |  | | | | | 0.03 | 0.004 | 6.77 | 1.43x10^-10^ |
|  |  |  | Age^2^ |  | | | | | -0.03 | 0.01 | -4.96 | 2.39x10^-6^ |
|  | Cubic | F | Model | 7.10 | 3 | 422 | 2.72x10^-4^ | 0.05 |  | | | |
|  |  |  | Age |  | | | | | 0.01 | 0.01 | 0.76 | .491 |
|  |  |  | Age^2^ |  | | | | | -0.02 | 0.01 | -3.59 | 7.25x10^-4^ |
|  |  |  | Age^3^ |  | | | | | 0.003 | 0.01 | 0.58 | .598 |
|  |  | M | Model | 29.72 | 3 | 409 | 9.84x10^-17^ | 0.18 |  | | | |
|  |  |  | Age |  | | | | | -0.01 | 0.01 | -1.45 | .178 |
|  |  |  | Age^2^ |  | | | | | -0.03 | 0.01 | -5.99 | 1.32x10^-8^ |
|  |  |  | Age^3^ |  | | | | | 0.03 | 0.01 | 4.58 | 1.27x10^-5^ |
|  | Log | F | Model | 13.31 | 1 | 424 | 6.22x10^-4^ | 0.03 |  | | | |
|  |  |  | Age |  | | | | | 0.03 | 0.01 | 3.65 | 6.22x10^-4^ |
|  |  | M | Model | 55.88 | 1 | 411 | 1.64x10^-12^ | 0.12 |  | | | |
|  |  |  | Age |  | | | | | 0.06 | 0.01 | 7.48 | 1.64x10^-12^ |
|  | S-curve | F | Model | 19.77 | 1 | 424 | 3.31x10^-5^ | 0.05 |  | | | |
|  |  |  | Age |  | | | | | -0.96 | 0.22 | -4.45 | 3.31x10^-5^ |
|  |  | M | Model | 78.52 | 1 | 411 | 1.05x10^-16^ | 0.16 |  | | | |
|  |  |  | Age |  | | | | | -2.11 | 0.24 | -8.86 | 1.05x10^-16^ |
| SNpc | Linear | F | Model | 5.71 | 1 | 423 | .025 | 0.01 |  | | | |
|  |  |  | Age |  | | | | | 0.001 | 2x10^-4^ | 2.39 | .025 |
|  |  | M | Model | 10.73 | 1 | 402 | .002 | 0.03 |  | | | |
|  |  |  | Age |  | | | | | 0.001 | 2x10^-4^ | 3.28 | .002 |
|  | Quadratic | F | Model | 4.62 | 2 | 422 | .016 | 0.02 |  | | | |
|  |  |  | Age |  | | | | | 0.01 | 0.004 | 2.53 | .018 |
|  |  |  | Age^2^ |  | | | | | -0.01 | 0.01 | -1.87 | .082 |
|  |  | M | Model | 16.48 | 2 | 401 | 3.30x10^-7^ | 0.08 |  | | | |
|  |  |  | Age |  | | | | | 0.02 | 0.004 | 3.80 | 2.90x10^-4^ |
|  |  |  | Age^2^ |  | | | | | -0.02 | 0.01 | -4.66 | 9.16x10^-6^ |
|  | Cubic | F | Model | 3.10 | 3 | 421 | .036 | 0.02 |  | | | |
|  |  |  | Age |  | | | | | 0.01 | 0.01 | 1.30 | .243 |
|  |  |  | Age^2^ |  | | | | | -0.01 | 0.01 | -1.74 | .107 |
|  |  |  | Age^3^ |  | | | | | -0.002 | 0.01 | -0.31 | .765 |
|  |  | M | Model | 14.07 | 3 | 400 | 2.64x10^-8^ | 0.10 |  | | | |
|  |  |  | Age |  | | | | | -0.01 | 0.01 | -1.18 | .278 |
|  |  |  | Age^2^ |  | | | | | -0.03 | 0.01 | -5.25 | 5.86x10^-7^ |
|  |  |  | Age^3^ |  | | | | | 0.02 | 0.01 | 2.94 | .005 |
|  | Log | F | Model | 7.61 | 1 | 423 | .009 | 0.02 |  | | | |
|  |  |  | Age |  | | | | | 0.02 | 0.01 | 2.76 | .009 |
|  |  | M | Model | 19.47 | 1 | 402 | 2.43x10^-5^ | 0.05 |  | | | |
|  |  |  | Age |  | | | | | 0.04 | 0.01 | 4.41 | 2.43x10^-5^ |
|  | S-curve | F | Model | 9.53 | 1 | 423 | .004 | 0.0220 |  | | | |
|  |  |  | Age |  | | | | | -0.79 | 0.26 | -3.09 | .004 |
|  |  | M | Model | 32.83 | 1 | 402 | 5.05x10^-8^ | 0.08 |  | | | |
|  |  |  | Age |  | | | | | -1.41 | 0.25 | -5.73 | 5.05x10^-8^ |
| SNpr | Linear | F | Model | 17.16 | 1 | 420 | 1.04x10^-4^ | 0.04 |  | | | |
|  |  |  | Age |  | | | | | 0.001 | 2x10^-4^ | 4.14 | 1.04x10^-4^ |
|  |  | M | Model | 37.65 | 1 | 405 | 5.84x10^-9^ | 0.09 |  | | | |
|  |  |  | Age |  | | | | | 0.002 | 2x10^-4^ | 6.14 | 5.84x10^-9^ |
|  | Quadratic | F | Model | 9.32 | 2 | 419 | 2.64x10^-4^ | 0.04 |  | | | |
|  |  |  | Age |  | | | | | 0.02 | 0.01 | 4.25 | 7.11x10^-5^ |
|  |  |  | Age^2^ |  | | | | | -0.007 | 0.01 | -1.21 | .279 |
|  |  | M | Model | 32.64 | 2 | 404 | 2.76x10^-13^ | 0.14 |  | | | |
|  |  |  | Age |  | | | | | 0.04 | 0.01 | 6.85 | 8.84x10^-11^ |
|  |  |  | Age^2^ |  | | | | | -0.03 | 0.01 | -5.04 | 1.66x10^-6^ |
|  | Cubic | F | Model | 6.20 | 3 | 418 | 7.52x10^-4^ | 0.04 |  | | | |
|  |  |  | Age |  | | | | | 0.02 | 0.01 | 1.70 | .114 |
|  |  |  | Age^2^ |  | | | | | -0.01 | 0.01 | -1.16 | .299 |
|  |  |  | Age^3^ |  | | | | | -1x10^-4^ | 0.01 | -0.02 | .984 |
|  |  | M | Model | 25.74 | 3 | 403 | 1.17x10^-14^ | 0.16 |  | | | |
|  |  |  | Age |  | | | | | -0.003 | 0.01 | -0.21 | .855 |
|  |  |  | Age^2^ |  | | | | | -0.04 | 0.01 | -5.76 | 4.54x10^-8^ |
|  |  |  | Age^3^ |  | | | | | 0.02 | 0.01 | 3.23 | .002 |
|  | Log | F | Model | 19.24 | 1 | 420 | 4.13x10^-5^ | 0.04 |  | | | |
|  |  |  | Age |  | | | | | 0.04 | 0.01 | 4.39 | 4.13x10^-5^ |
|  |  | M | Model | 54.32 | 1 | 405 | 3.23x10^-12^ | 0.12 |  | | | |
|  |  |  | Age |  | | | | | 0.07 | 0.01 | 7.37 | 3.23x10^-12^ |
|  | S-curve | F | Model | 22.18 | 1 | 420 | 1.05x10^-5^ | 0.05 |  | | | |
|  |  |  | Age |  | | | | | -1.29 | 0.27 | -4.71 | 1.05x10^-5^ |
|  |  | M | Model | 76.92 | 1 | 405 | 2.03x10^-16^ | 0.16 |  | | | |
|  |  |  | Age |  | | | | | -2.40 | 0.27 | -8.77 | 2.03x10^-16^ |
| Thalamus | Linear | F | Model | 13.66 | 1 | 430 | 5.45x10^-4^ | 0.03 |  | | | |
|  |  |  | Age |  | | | | | -2x10^-4^ | 5.5x10^-5^ | -3.70 | 5.45x10^-4^ |
|  |  | M | Model | 0.46 | 1 | 413 | .543 | 0.001 |  | | | |
|  |  |  | Age |  | | | | | 3.63x10^-5^ | 5.35x10^-5^ | 0.68 | .543 |
|  | Quadratic | F | Model | 13.32 | 2 | 429 | 7.93x10^-6^ | 0.06 |  | | | |
|  |  |  | Age |  | | | | | -0.005 | 0.001 | -4.12 | 1.12x10^-4^ |
|  |  |  | Age^2^ |  | | | | | -0.004 | 0.001 | -3.09 | 3.59x10^-3^ |
|  |  | M | Model | 4.43 | 2 | 412 | .018 | 0.02 |  | | | |
|  |  |  | Age |  | | | | | 3x10^-4^ | 0.001 | 0.29 | .818 |
|  |  |  | Age^2^ |  | | | | | -0.004 | 0.001 | -2.98 | .005 |
|  | Cubic | F | Model | 13.60 | 3 | 428 | 6.84x10^-8^ | 0.09 |  | | | |
|  |  |  | Age |  | | | | | -0.02 | 0.003 | -4.95 | 3.69x10^-6^ |
|  |  |  | Age^2^ |  | | | | | -0.01 | 0.001 | -3.80 | 3.78x10^-4^ |
|  |  |  | Age^3^ |  | | | | | 0.01 | 0.002 | 3.59 | 7.25x10^-4^ |
|  |  | M | Model | 9.92 | 3 | 411 | 5.35x10^-6^ | 0.07 |  | | | |
|  |  |  | Age |  | | | | | -0.01 | 0.003 | -4.09 | 9.24x10^-5^ |
|  |  |  | Age^2^ |  | | | | | -0.005 | 0.001 | -3.93 | 1.74x10^-4^ |
|  |  |  | Age^3^ |  | | | | | 0.01 | 0.002 | 4.50 | 1.74x10^-5^ |
|  | Log | F | Model | 5.52 | 1 | 430 | .027 | 0.01 |  | | | |
|  |  |  | Age |  | | | | | -0.01 | 0.002 | -2.35 | .027 |
|  |  | M | Model | 4.48 | 1 | 413 | .047 | 0.01 |  | | | |
|  |  |  | Age |  | | | | | 0.004 | 0.002 | 2.12 | .047 |
|  | S-curve | F | Model | 0.58 | 1 | 430 | .491 | 0.001 |  | | | |
|  |  |  | Age |  | | | | | 0.05 | 0.06 | 0.76 | .491 |
|  |  | M | Model | 9.98 | 1 | 413 | .003 | 0.02 |  | | | |
|  |  |  | Age |  | | | | | -0.19 | 0.06 | -3.16 | .003 |
| VTA | Linear | F | Model | 61.46 | 1 | 409 | 2.06x10^-13^ | 0.13 |  | | | |
|  |  |  | Age |  | | | | | 0.002 | 3x10^-4^ | 7.84 | 2.06x10^-13^ |
|  |  | M | Model | 82.23 | 1 | 397 | 2.72x10^-17^ | 0.17 |  | | | |
|  |  |  | Age |  | | | | | 0.003 | 3x10^-4^ | 9.07 | 2.72x10^-17^ |
|  | Quadratic | F | Model | 57.12 | 2 | 408 | 4.39x10^-21^ | 0.22 |  | | | |
|  |  |  | Age |  | | | | | 0.05 | 0.01 | 8.85 | 2.61x10^-16^ |
|  |  |  | Age^2^ |  | | | | | -0.04 | 0.01 | -6.78 | 1.74x10^-10^ |
|  |  | M | Model | 51.42 | 2 | 396 | 7.97x10^-20^ | 0.21 |  | | | |
|  |  |  | Age |  | | | | | 0.06 | 0.01 | 9.67 | 3.00x10^-19^ |
|  |  |  | Age^2^ |  | | | | | -0.03 | 0.01 | -4.15 | 7.28x10^-5^ |
|  | Cubic | F | Model | 39.97 | 3 | 407 | 4.39x10^-21^ | 0.23 |  | | | |
|  |  |  | Age |  | | | | | 0.02 | 0.01 | 1.61 | .136 |
|  |  |  | Age^2^ |  | | | | | -0.05 | 0.01 | -7.14 | 2.08x10^-11^ |
|  |  |  | Age^3^ |  | | | | | 0.02 | 0.01 | 2.16 | .042 |
|  |  | M | Model | 39.69 | 3 | 395 | 1.41x10^-21^ | 0.23 |  | | | |
|  |  |  | Age |  | | | | | 0.01 | 0.01 | 0.57 | .612 |
|  |  |  | Age^2^ |  | | | | | -0.03 | 0.01 | -4.95 | 2.47x10^-6^ |
|  |  |  | Age^3^ |  | | | | | 0.03 | 0.01 | 3.62 | 5.59x10^-4^ |
|  | Log | F | Model | 90.50 | 1 | 409 | 2.46x10^-18^ | 0.18 |  | | | |
|  |  |  | Age |  | | | | | 0.10 | 0.01 | 9.51 | 2.46x10^-18^ |
|  |  | M | Model | 104.73 | 1 | 397 | 3.70x10^-21^ | 0.21 |  | | | |
|  |  |  | Age |  | | | | | 0.11 | 0.01 | 10.23 | 3.70x10^-21^ |
|  | S-curve | F | Model | 139.63 | 1 | 409 | 4.15x10^-26^ | 0.26 |  | | | |
|  |  |  | Age |  | | | | | -3.82 | 0.32 | -11.82 | 4.15x10^-26^ |
|  |  | M | Model | 146.93 | 1 | 397 | 5.47x10^-28^ | 0.27 |  | | | |
|  |  |  | Age |  | | | | | -4.18 | 0.35 | -12.12 | 5.47x10^-28^ |

*Note.* Table includes models of five curves tested for each of the 10 ROIs within female and male participants separately.

Supplementary Table S2: AIC Comparisons for 10 Major ROIs

| Region | Curve | Female Models | | Male Models | |
| --- | --- | --- | --- | --- | --- |
|  |  | Original | Weighted | Original | Weighted |
| Caudate | | | | | |
|  | Linear | -1875 | - | - | n.s. |
|  | Quadratic | -1896.42 | - | - | **-1758.75** |
|  | Cubic | **-1906.68** | - | - | n.s. |
|  | Log | -1861.13 | - | - | n.s. |
|  | S-Curve | -1694.26 | - | - | n.s. |
| Hippocampus | | | | | |
|  | Linear | n.s. | - | n.s. | - |
|  | Quadratic | n.s. | - | **-1528.83** | - |
|  | Cubic | n.s. | - | n.s. | - |
|  | Log | n.s. | - | n.s. | - |
|  | S-Curve | n.s. | - | n.s. | - |
| Nucleus Accumbens | | | | | |
|  | Linear | - | **-1044.62** | -915.879 | - |
|  | Quadratic | - | -1041.86 | n.s. | - |
|  | Cubic | - | n.s. | n.s. | - |
|  | Log | - | -1043.31 | **-918.17** | - |
|  | S-Curve | - | -996.24 | -878.085 | - |
| Pallidum | | | | | |
|  | Linear | - | -716.093 | - | -641.73 |
|  | Quadratic | - | -728.193 | - | -640.33 |
|  | Cubic | - | n.s. | - | -633.95 |
|  | Log | - | -727.837 | - | -636.47 |
|  | S-Curve | - | **-913.378** | - | **-798.2** |
| Putamen | | | | | |
|  | Linear | - | -1340.25 | - | **-1237.09** |
|  | Quadratic | - | -1339.53 | - | -1235.13 |
|  | Cubic | - | -1336.22 | - | -1233.17 |
|  | Log | - | **-1344.81** | - | -1214.2 |
|  | S-Curve | - | -1324.69 | - | -1170.51 |
| Red Nucleus | | | | | |
|  | Linear | -953.474 | - | -820.335 | - |
|  | Quadratic | -964.114 | - | -842.368 | - |
|  | Cubic | -962.448 | - | -861.004 | - |
|  | Log | -958.633 | - | -836.213 | - |
|  | S-Curve | **-1013.76** | - | **-920.759** | - |
| Substantia Nigra pars compacta | | | | | |
|  | Linear | -831.773 | - | -821.008 | - |
|  | Quadratic | -833.274 | - | -840.287 | - |
|  | Cubic | -831.374 | - | -846.903 | - |
|  | Log | -833.651 | - | -829.479 | - |
|  | S-Curve | **-857.523** | - | **-875.274** | - |
| Substantia Nigra pars reticulata | | | | | |
|  | Linear | -732.501 | - | -676.513 | - |
|  | Quadratic | -731.971 | - | -699.299 | - |
|  | Cubic | -729.971 | - | -707.685 | - |
|  | Log | -734.511 | - | -691.557 | - |
|  | S-Curve | **-806.375** | - | **-797.317** | - |
| Ventral Tegmental Area | | | | | |
|  | Linear | -622.68 | - | -575.033 | - |
|  | Quadratic | -664.6 | - | -590.038 | - |
|  | Cubic | -667.279 | - | -601.047 | - |
|  | Log | -647.298 | - | -593.34 | - |
|  | S-Curve | **-670.91** | - | **-612.977** | - |
| Thalamus | | | | | |
|  | Linear | - | -1972.49 | - | -1927.25 |
|  | Quadratic | - | -1971.51 | - | -1920.23 |
|  | Cubic | - | **-1980.01** | - | **-1942.95** |
|  | Log | - | -1970.9 | - | -1934.25 |
|  | S-Curve | - | -1850 | - | -1819.18 |

*Note.* Lowest AIC values are shown in bold font. n.s. = not significant either before or after multiple comparison correction.

Supplementary Table S3: All Tested Regressions for 14 Thalamic Nuclei

| ROI | Slope | Sex | Variable | *F* | df1 | df2 | *p_FDR_* | *R^2^* | *b* | SE | *T* | *p_FDR_* |
| --- | --- | --- | --- | --- | --- | --- | --- | --- | --- | --- | --- | --- |
| AV | Linear | F | Model | 1.76 | 1 | 442 | .281 | 0.004 |  |  |  |  |
|  |  |  | Age |  |  |  |  |  | 2.00x10^-4^ | 1.00x10^-4^ | 1.33 | .281 |
|  |  | M | Model | 7.09 | 1 | 415 | .014 | 0.017 |  |  |  |  |
|  |  |  | Age |  |  |  |  |  | 3.00x10^-4^ | 1.00x10^-4^ | 2.66 | .014 |
|  | Quadratic | F | Model | 20.13 | 2 | 441 | 1.56x10^-7^ | 0.084 |  |  |  |  |
|  |  |  | Age |  |  |  |  |  | 0.003 | 0.003 | 1.25 | .309 |
|  |  |  | Age^2^ |  |  |  |  |  | -0.02 | 0.003 | -6.33 | 6.11x10^-8^ |
|  |  | M | Model | 12.15 | 2 | 414 | 2.66x10^-5^ | 0.06 |  |  |  |  |
|  |  |  | Age |  |  |  |  |  | 0.01 | 0.003 | 3.43 | .002 |
|  |  |  | Age^2^ |  |  |  |  |  | -0.01 | 0.003 | -3.34 | .002 |
|  | Cubic | F | Model | 13.41 | 3 | 440 | 5.61x10^-7^ | 0.08 |  |  |  |  |
|  |  |  | Age |  |  |  |  |  | -0.003 | 0.01 | -0.49 | .784 |
|  |  |  | Age^2^ |  |  |  |  |  | -0.02 | 0.003 | -6.28 | 6.11x10^-8^ |
|  |  |  | Age^3^ |  |  |  |  |  | 0.004 | 0.003 | 1.04 | .407 |
|  |  | M | Model | 13.78 | 3 | 413 | 8.15x10^-8^ | 0.09 |  |  |  |  |
|  |  |  | Age |  |  |  |  |  | -0.02 | 0.01 | -2.38 | .028 |
|  |  |  | Age^2^ |  |  |  |  |  | -0.01 | 0.003 | -3.29 | .002 |
|  |  |  | Age^3^ |  |  |  |  |  | 0.02 | 0.004 | 3.82 | 4.53x10^-4^ |
|  | Log | F | Model | 9.87 | 1 | 442 | .009 | 0.02 |  |  |  |  |
|  |  |  | Age |  |  |  |  |  | 0.02 | 0.01 | 3.14 | .009 |
|  |  | M | Model | 16.84 | 1 | 415 | 1.57x10-4 | 0.04 |  |  |  |  |
|  |  |  | Age |  |  |  |  |  | 0.02 | 0.01 | 4.10 | 1.57x10^-4^ |
|  | S-curve | F | Model | 22.04 | 1 | 442 | 4.75x10^-5^ | 0.05 |  |  |  |  |
|  |  |  | Age |  |  |  |  |  | -0.65 | 0.14 | -4.69 | 4.75x10^-5^ |
|  |  | M | Model | 30.29 | 1 | 415 | 3.04x10^-7^ | 0.07 |  |  |  |  |
|  |  |  | Age |  |  |  |  |  | -0.77 | 0.14 | -5.50 | 3.04x10^-7^ |
| IL | Linear | F | Model | 3.15 | 1 | 431 | .141 | 0.01 |  |  |  |  |
|  |  |  | Age |  |  |  |  |  | -1.00x10^-4^ | 1.00x10^-4^ | -1.78 | .141 |
|  |  | M | Model | 10.57 | 1 | 403 | .003 | 0.03 |  |  |  |  |
|  |  |  | Age |  |  |  |  |  | 2.00x10^-4^ | 1.00x10^-4^ | 3.25 | .003 |
|  | Quadratic | F | Model | 1.11 | 2 | 430 | .447 | 0.01 |  |  |  |  |
|  |  |  | Age |  |  |  |  |  | -0.002 | 0.001 | -1.39 | .265 |
|  |  |  | Age^2^ |  |  |  |  |  | 3.00x10^-4^ | 0.001 | 0.20 | .950 |
|  |  | M | Model | 7.30 | 2 | 402 | .002 | 0.04 |  |  |  |  |
|  |  |  | Age |  |  |  |  |  | 0.004 | 0.001 | 3.11 | .004 |
|  |  |  | Age^2^ |  |  |  |  |  | -0.002 | 0.002 | -1.35 | .226 |
|  | Cubic | F | Model | 3.56 | 3 | 429 | .044 | 0.02 |  |  |  |  |
|  |  |  | Age |  |  |  |  |  | -0.01 | 0.003 | -3.16 | .009 |
|  |  |  | Age^2^ |  |  |  |  |  | 1.00x10^-4^ | 0.002 | 0.07 | .965 |
|  |  |  | Age^3^ |  |  |  |  |  | 0.01 | 0.002 | 2.71 | .025 |
|  |  | M | Model | 5.44 | 3 | 401 | .003 | 0.04 |  |  |  |  |
|  |  |  | Age |  |  |  |  |  | -0.004 | 0.003 | -1.25 | .263 |
|  |  |  | Age^2^ |  |  |  |  |  | -0.001 | 0.002 | -0.64 | .573 |
|  |  |  | Age^3^ |  |  |  |  |  | 0.004 | 0.002 | 2.44 | .024 |
|  | Log | F | Model | 2.15 | 1 | 431 | .231 | 0.01 |  |  |  |  |
|  |  |  | Age |  |  |  |  |  | -0.003 | 0.002 | -1.46 | .231 |
|  |  | M | Model | 14.53 | 1 | 403 | 4.53x10^-4^ | 0.04 |  |  |  |  |
|  |  |  | Age |  |  |  |  |  | 0.01 | 0.002 | 3.81 | 4.53x10^-4^ |
|  | S-curve | F | Model | 3.04 | 1 | 431 | .144 | 0.01 |  |  |  |  |
|  |  |  | Age |  |  |  |  |  | 0.09 | 0.05 | 1.74 | .144 |
|  |  | M | Model | 12.70 | 1 | 403 | .001 | 0.03 |  |  |  |  |
|  |  |  | Age |  |  |  |  |  | -0.22 | 0.06 | -3.56 | .001 |
| LGN | Linear | F | Model | 3.64 | 1 | 439 | .114 | 0.01 |  |  |  |  |
|  |  |  | Age |  |  |  |  |  | 1.00x10^-4^ | 1.00x10^-4^ | 1.91 | .114 |
|  |  | M | Model | 32.72 | 1 | 421 | 1.08x10^-7^ | 0.07 |  |  |  |  |
|  |  |  | Age |  |  |  |  |  | 4.00x10^-4^ | 1.00x10^-4^ | 5.72 | 1.08x10^-7^ |
|  | Quadratic | F | Model | 2.54 | 2 | 438 | .144 | 0.01 |  |  |  |  |
|  |  |  | Age |  |  |  |  |  | 0.003 | 0.002 | 2.00 | .097 |
|  |  |  | Age^2^ |  |  |  |  |  | -0.002 | 0.002 | -0.92 | .472 |
|  |  | M | Model | 31.14 | 2 | 420 | 1.12x10^-11^ | 0.13 |  |  |  |  |
|  |  |  | Age |  |  |  |  |  | 0.01 | 0.001 | 6.12 | 1.50x10^-8^ |
|  |  |  | Age^2^ |  |  |  |  |  | -0.01 | 0.002 | -4.72 | 1.32x10^-5^ |
|  | Cubic | F | Model | 4.49 | 3 | 437 | .017 | 0.03 |  |  |  |  |
|  |  |  | Age |  |  |  |  |  | -0.004 | 0.004 | -1.10 | .385 |
|  |  |  | Age^2^ |  |  |  |  |  | -0.002 | 0.002 | -1.26 | .309 |
|  |  |  | Age^3^ |  |  |  |  |  | 0.004 | 0.002 | 2.18 | .069 |
|  |  | M | Model | 23.25 | 3 | 419 | 3.72x10^-12^ | 0.14 |  |  |  |  |
|  |  |  | Age |  |  |  |  |  | 0.01 | 0.004 | 1.22 | .273 |
|  |  |  | Age^2^ |  |  |  |  |  | -0.01 | 0.002 | -4.88 | 6.47x10^-6^ |
|  |  |  | Age^3^ |  |  |  |  |  | 0.002 | 0.002 | 1.12 | .316 |
|  | Log | F | Model | 5.93 | 1 | 439 | .044 | 0.01 |  |  |  |  |
|  |  |  | Age |  |  |  |  |  | 0.01 | 0.003 | 2.44 | .0441 |
|  |  | M | Model | 51.52 | 1 | 421 | 4.52x10^-11^ | 0.11 |  |  |  |  |
|  |  |  | Age |  |  |  |  |  | 0.02 | 0.003 | 7.18 | 4.52x10^-11^ |
|  | S-curve | F | Model | 10.31 | 1 | 439 | .008 | 0.02 |  |  |  |  |
|  |  |  | Age |  |  |  |  |  | -0.26 | 0.08 | -3.21 | .008 |
|  |  | M | Model | 67.97 | 1 | 421 | 1.94x10^-13^ | 0.14 |  |  |  |  |
|  |  |  | Age |  |  |  |  |  | -0.62 | 0.08 | -8.24 | 1.94x10^-13^ |
| LP | Linear | F | Model | 5.12 | 1 | 436 | .058 | 0.01 |  |  |  |  |
|  |  |  | Age |  |  |  |  |  | -2.00x10^-4^ | 1.00x10^-4^ | -2.26 | .058 |
|  |  | M | Model | 0.04 | 1 | 409 | .879 | 1x10^-4^ |  |  |  |  |
|  |  |  | Age |  |  |  |  |  | -2.29x10^-5^ | 1.00x10^-4^ | -0.19 | .879 |
|  | Quadratic | F | Model | 4.30 | 2 | 435 | .044 | 0.02 |  |  |  |  |
|  |  |  | Age |  |  |  |  |  | -0.01 | 0.002 | -2.26 | .058 |
|  |  |  | Age^2^ |  |  |  |  |  | -0.01 | 0.003 | -1.88 | .118 |
|  |  | M | Model | 5.52 | 2 | 408 | .008 | 0.03 |  |  |  |  |
|  |  |  | Age |  |  |  |  |  | -0.003 | 0.003 | -0.89 | .428 |
|  |  |  | Age^2^ |  |  |  |  |  | -0.01 | 0.003 | -3.17 | .003 |
|  | Cubic | F | Model | 2.87 | 3 | 434 | .082 | 0.02 |  |  |  |  |
|  |  |  | Age |  |  |  |  |  | -0.01 | 0.01 | -1.09 | .388 |
|  |  |  | Age^2^ |  |  |  |  |  | -0.01 | 0.003 | -1.88 | .118 |
|  |  |  | Age^3^ |  |  |  |  |  | 0.001 | 0.003 | 0.17 | .954 |
|  |  | M | Model | 5.62 | 3 | 407 | .002 | 0.04 |  |  |  |  |
|  |  |  | Age |  |  |  |  |  | -0.02 | 0.01 | -2.48 | .023 |
|  |  |  | Age^2^ |  |  |  |  |  | -0.01 | 0.003 | -3.63 | 8.20x10^-4^ |
|  |  |  | Age^3^ |  |  |  |  |  | 0.01 | 0.004 | 2.39 | .027 |
|  | Log | F | Model | 3.25 | 1 | 436 | .138 | 0.01 |  |  |  |  |
|  |  |  | Age |  |  |  |  |  | -0.01 | 0.004 | -1.80 | .138 |
|  |  | M | Model | 0.60 | 1 | 409 | .494 | 0.001 |  |  |  |  |
|  |  |  | Age |  |  |  |  |  | 0.004 | 0.01 | 0.77 | .494 |
|  | S-curve | F | Model | 1.90 | 1 | 436 | .2660 | 0.004 |  |  |  |  |
|  |  |  | Age |  |  |  |  |  | 0.19 | 0.13 | 1.38 | .266 |
|  |  | M | Model | 2.20 | 1 | 409 | .185 | 0.01 |  |  |  |  |
|  |  |  | Age |  |  |  |  |  | -0.22 | 0.15 | -1.48 | .185 |
| MDl | Linear | F | Model | 10.91 | 1 | 434 | .006 | 0.03 |  |  |  |  |
|  |  |  | Age |  |  |  |  |  | -2.00x10^-4^ | 1.00x10^-4^ | -3.30 | .006 |
|  |  | M | Model | 2.22 | 1 | 414 | .185 | 0.01 |  |  |  |  |
|  |  |  | Age |  |  |  |  |  | 1.00x10^-4^ | 1.00x10^-4^ | 1.49 | .185 |
|  | Quadratic | F | Model | 15.04 | 2 | 433 | 9.80x10^-6^ | 0.07 |  |  |  |  |
|  |  |  | Age |  |  |  |  |  | -0.01 | 0.002 | -4.05 | 5.55x10^-4^ |
|  |  |  | Age^2^ |  |  |  |  |  | -0.01 | 0.002 | -3.92 | 8.18x10^-4^ |
|  |  | M | Model | 7.32 | 2 | 413 | .002 | 0.03 |  |  |  |  |
|  |  |  | Age |  |  |  |  |  | 0.002 | 0.001 | 1.17 | .299 |
|  |  |  | Age^2^ |  |  |  |  |  | -0.0060 | 0.002 | -3.75 | 5.36x10^-4^ |
|  | Cubic | F | Model | 15.51 | 3 | 432 | 6.11x10^-8^ | 0.10 |  |  |  |  |
|  |  |  | Age |  |  |  |  |  | -0.02 | 0.003 | -5.17 | 8.31x10^-6^ |
|  |  |  | Age^2^ |  |  |  |  |  | -0.01 | 0.002 | -4.53 | 8.33x10^-5^ |
|  |  |  | Age^3^ |  |  |  |  |  | 0.01 | 0.002 | 3.88 | 9.24x10^-4^ |
|  |  | M | Model | 12.54 | 3 | 412 | 3.35x10^-7^ | 0.08 |  |  |  |  |
|  |  |  | Age |  |  |  |  |  | -0.01 | 0.003 | -3.79 | 4.86x10^-4^ |
|  |  |  | Age^2^ |  |  |  |  |  | -0.01 | 0.002 | -4.79 | 9.79x10^-6^ |
|  |  |  | Age^3^ |  |  |  |  |  | 0.01 | 0.002 | 4.56 | 2.47x10^-5^ |
|  | Log | F | Model | 3.22 | 1 | 434 | .138 | 0.01 |  |  |  |  |
|  |  |  | Age |  |  |  |  |  | -0.004 | 0.002 | -1.79 | .138 |
|  |  | M | Model | 10.24 | 1 | 414 | .003 | 0.02 |  |  |  |  |
|  |  |  | Age |  |  |  |  |  | 0.01 | 0.002 | 3.20 | .003 |
|  | S-curve | F | Model | 0.03 | 1 | 434 | .953 | 1.00x10^-4^ |  |  |  |  |
|  |  |  | Age |  |  |  |  |  | -0.01 | 0.08 | -0.18 | .953 |
|  |  | M | Model | 22.15 | 1 | 414 | 1.33x10^-5^ | 0.05 |  |  |  |  |
|  |  |  | Age |  |  |  |  |  | -0.36 | 0.08 | -4.71 | 1.33x10^-5^ |
| MDm | Linear | F | Model | 9.89 | 1 | 432 | .009 | 0.02 |  |  |  |  |
|  |  |  | Age |  |  |  |  |  | -2.00x10^-4^ | 1.00x10^-4^ | -3.14 | .009 |
|  |  | M | Model | 0.22 | 1 | 415 | .688 | 0.001 |  |  |  |  |
|  |  |  | Age |  |  |  |  |  | -3.00x10^-5^ | 1.00x10^-4^ | -0.46 | .688 |
|  | Quadratic | F | Model | 12.31 | 2 | 431 | 7.30x10^-5^ | 0.05 |  |  |  |  |
|  |  |  | Age |  |  |  |  |  | -0.006 | 0.002 | -3.94 | 8.08x10^-4^ |
|  |  |  | Age^2^ |  |  |  |  |  | -0.01 | 0.002 | -3.25 | .007 |
|  |  | M | Model | 5.87 | 2 | 414 | .006 | 0.03 |  |  |  |  |
|  |  |  | Age |  |  |  |  |  | -0.002 | 0.002 | -1.60 | .155 |
|  |  |  | Age^2^ |  |  |  |  |  | -0.005 | 0.002 | -2.98 | .006 |
|  | Cubic | F | Model | 15.64 | 3 | 430 | 6.11x10^-8^ | 0.10 |  |  |  |  |
|  |  |  | Age |  |  |  |  |  | -0.02 | 0.004 | -5.88 | 2.49x10^-7^ |
|  |  |  | Age^2^ |  |  |  |  |  | -0.01 | 0.002 | -3.93 | 8.08x10^-4^ |
|  |  |  | Age^3^ |  |  |  |  |  | 0.01 | 0.002 | 4.74 | 4.75x10^-5^ |
|  |  | M | Model | 10.30 | 3 | 413 | 6.47x10^-6^ | 0.07 |  |  |  |  |
|  |  |  | Age |  |  |  |  |  | -0.02 | 0.004 | -4.72 | 1.32x10^-5^ |
|  |  |  | Age^2^ |  |  |  |  |  | -0.01 | 0.002 | -3.77 | 5.08x10^-4^ |
|  |  |  | Age^3^ |  |  |  |  |  | 0.01 | 0.002 | 4.56 | 2.47x10^-5^ |
|  | Log | F | Model | 2.53 | 1 | 432 | .189 | 0.01 |  |  |  |  |
|  |  |  | Age |  |  |  |  |  | -0.004 | 0.003 | -1.59 | .189 |
|  |  | M | Model | 1.26 | 1 | 415 | .316 | 0.003 |  |  |  |  |
|  |  |  | Age |  |  |  |  |  | 0.003 | 0.002 | 1.12 | .316 |
|  | S-curve | F | Model | 0.23 | 1 | 432 | .784 | 0.001 |  |  |  |  |
|  |  |  | Age |  |  |  |  |  | -0.04 | 0.08 | -0.48 | .784 |
|  |  | M | Model | 6.59 | 1 | 415 | .018 | 0.02 |  |  |  |  |
|  |  |  | Age |  |  |  |  |  | -0.17 | 0.07 | -2.57 | .018 |
| MGN | Linear | F | Model | 0.11 | 1 | 432 | .876 | 3.00x10^-4^ |  |  |  |  |
|  |  |  | Age |  |  |  |  |  | -4.20x10^-5^ | 1.00x10^-4^ | -0.33 | .876 |
|  |  | M | Model | 1.83 | 1 | 414 | .226 | 0.004 |  |  |  |  |
|  |  |  | Age |  |  |  |  |  | 2.00x10^-4^ | 1.00x10^-4^ | 1.35 | .226 |
|  | Quadratic | F | Model | 0.07 | 2 | 431 | .958 | 3.29x10^-4^ |  |  |  |  |
|  |  |  | Age |  |  |  |  |  | -0.001 | 0.003 | -0.34 | .876 |
|  |  |  | Age^2^ |  |  |  |  |  | -0.001 | 0.003 | -0.15 | .957 |
|  |  | M | Model | 10.35 | 2 | 413 | 1.36x10^-4^ | 0.05 |  |  |  |  |
|  |  |  | Age |  |  |  |  |  | 0.003 | 0.003 | 1.08 | .330 |
|  |  |  | Age^2^ |  |  |  |  |  | -0.01 | 0.003 | -4.50 | 3.15x10^-5^ |
|  | Cubic | F | Model | 0.02 | 3 | 430 | .997 | 1.25x10^-4^ |  |  |  |  |
|  |  |  | Age |  |  |  |  |  | -7.00x10^-4^ | 0.01 | -0.10 | .957 |
|  |  |  | Age^2^ |  |  |  |  |  | -4.00x10^-4^ | 0.003 | -0.13 | .957 |
|  |  |  | Age^3^ |  |  |  |  |  | 1.00x10^-4^ | 0.004 | 0.02 | .992 |
|  |  | M | Model | 6.94 | 3 | 412 | 4.33x10^-4^ | 0.05 |  |  |  |  |
|  |  |  | Age |  |  |  |  |  | 0.01 | 0.01 | 0.79 | .489 |
|  |  |  | Age^2^ |  |  |  |  |  | -0.01 | 0.003 | -4.28 | 8.14x10^-5^ |
|  |  |  | Age^3^ |  |  |  |  |  | -0.002 | 0.004 | -0.44 | .700 |
|  | Log | F | Model | 0.02 | 1 | 432 | .957 | 3.55x10^-5^ |  |  |  |  |
|  |  |  | Age |  |  |  |  |  | -0.001 | 0.01 | -0.12 | .957 |
|  |  | M | Model | 5.99 | 1 | 414 | .024 | 0.01 |  |  |  |  |
|  |  |  | Age |  |  |  |  |  | 0.01 | 0.01 | 2.45 | .024 |
|  | S-curve | F | Model | 0.06 | 1 | 432 | .936 | 1.30x10^-4^ |  |  |  |  |
|  |  |  | Age |  |  |  |  |  | -0.03 | 0.12 | -0.24 | .936 |
|  |  | M | Model | 10.56 | 1 | 414 | .003 | 0.03 |  |  |  |  |
|  |  |  | Age |  |  |  |  |  | -0.45 | 0.14 | -3.25 | .003 |
| PuA | Linear | F | Model | 6.73 | 1 | 431 | .033 | 0.02 |  |  |  |  |
|  |  |  | Age |  |  |  |  |  | 2.00x10^-4^ | 1.00x10^-4^ | 2.59 | .033 |
|  |  | M | Model | 35.21 | 1 | 418 | 3.76x10^-8^ | 0.08 |  |  |  |  |
|  |  |  | Age |  |  |  |  |  | 0.001 | 1.00x10^-4^ | 5.93 | 3.76x10^-8^ |
|  | Quadratic | F | Model | 4.42 | 2 | 430 | .040 | 0.02 |  |  |  |  |
|  |  |  | Age |  |  |  |  |  | 0.01 | 0.002 | 2.66 | .028 |
|  |  |  | Age^2^ |  |  |  |  |  | -0.003 | 0.002 | -1.23 | .319 |
|  |  | M | Model | 27.01 | 2 | 417 | 1.13x10^-10^ | 0.12 |  |  |  |  |
|  |  |  | Age |  |  |  |  |  | 0.01 | 0.002 | 6.36 | 3.94x10^-9^ |
|  |  |  | Age^2^ |  |  |  |  |  | -0.01 | 0.002 | -4.17 | 1.22x10^-4^ |
|  | Cubic | F | Model | 3.45 | 3 | 429 | .046 | 0.02 |  |  |  |  |
|  |  |  | Age |  |  |  |  |  | 0.001 | 0.004 | 0.26 | .924 |
|  |  |  | Age^2^ |  |  |  |  |  | -0.003 | 0.002 | -1.37 | .268 |
|  |  |  | Age^3^ |  |  |  |  |  | 0.002 | 0.002 | 0.87 | .500 |
|  |  | M | Model | 19.69 | 3 | 416 | 7.74x10^-11^ | 0.12 |  |  |  |  |
|  |  |  | Age |  |  |  |  |  | 0.003 | 0.01 | 0.59 | .607 |
|  |  |  | Age^2^ |  |  |  |  |  | -0.01 | 0.002 | -4.56 | 2.50x10^-5^ |
|  |  |  | Age^3^ |  |  |  |  |  | 0.01 | 0.003 | 2.14 | .050 |
|  | Log | F | Model | 9.27 | 1 | 431 | .011 | 0.02 |  |  |  |  |
|  |  |  | Age |  |  |  |  |  | 0.01 | 0.003 | 3.04 | .011 |
|  |  | M | Model | 46.61 | 1 | 418 | 3.27x10^-10^ | 0.10 |  |  |  |  |
|  |  |  | Age |  |  |  |  |  | 0.02 | 0.004 | 6.83 | 3.27x10^-10^ |
|  | S-curve | F | Model | 12.32 | 1 | 431 | .003 | 0.03 |  |  |  |  |
|  |  |  | Age |  |  |  |  |  | -0.36 | 0.10 | -3.51 | .003 |
|  |  | M | Model | 53.71 | 1 | 418 | 2.75x10^-11^ | 0.11 |  |  |  |  |
|  |  |  | Age |  |  |  |  |  | -0.91 | 0.12 | -7.33 | 2.75x10^-11^ |
| PuI | Linear | F | Model | 6.52 | 1 | 441 | .036 | 0.02 |  |  |  |  |
|  |  |  | Age |  |  |  |  |  | -2.00x10^-4^ | 1.00x10^-4^ | -2.55 | .036 |
|  |  | M | Model | 0.07 | 1 | 415 | .828 | 1.77x10^-4^ |  |  |  |  |
|  |  |  | Age |  |  |  |  |  | 1.88x10^-5^ | 1.00x10^-4^ | 0.27 | .828 |
|  | Quadratic | F | Model | 4.19 | 2 | 440 | .045 | 0.02 |  |  |  |  |
|  |  |  | Age |  |  |  |  |  | -0.003 | 0.001 | -2.40 | .047 |
|  |  |  | Age^2^ |  |  |  |  |  | -0.002 | 0.002 | -1.36 | .268 |
|  |  | M | Model | 1.28 | 2 | 414 | .330 | 0.01 |  |  |  |  |
|  |  |  | Age |  |  |  |  |  | 0.001 | 0.002 | 0.54 | .645 |
|  |  |  | Age^2^ |  |  |  |  |  | -0.003 | 0.002 | -1.56 | .167 |
|  | Cubic | F | Model | 6.82 | 3 | 439 | .001 | 0.05 |  |  |  |  |
|  |  |  | Age |  |  |  |  |  | -0.02 | 0.004 | -4.13 | 4.15x10^-4^ |
|  |  |  | Age^2^ |  |  |  |  |  | -0.004 | 0.002 | -2.29 | .057 |
|  |  |  | Age^3^ |  |  |  |  |  | 0.01 | 0.002 | 3.45 | .004 |
|  |  | M | Model | 5.81 | 3 | 413 | .002 | 0.04 |  |  |  |  |
|  |  |  | Age |  |  |  |  |  | -0.01 | 0.004 | -3.38 | .002 |
|  |  |  | Age^2^ |  |  |  |  |  | -0.01 | 0.002 | -2.49 | .022 |
|  |  |  | Age^3^ |  |  |  |  |  | 0.01 | 0.002 | 3.81 | 4.56x10^-4^ |
|  | Log | F | Model | 3.87 | 1 | 441 | .103 | 0.01 |  |  |  |  |
|  |  |  | Age |  |  |  |  |  | -0.01 | 0.003 | -1.97 | .103 |
|  |  | M | Model | 0.96 | 1 | 415 | .379 | 0.002 |  |  |  |  |
|  |  |  | Age |  |  |  |  |  | 0.003 | 0.003 | 0.98 | .379 |
|  | S-curve | F | Model | 1.31 | 1 | 441 | .364 | 0.003 |  |  |  |  |
|  |  |  | Age |  |  |  |  |  | 0.12 | 0.10 | 1.14 | .364 |
|  |  | M | Model | 2.25 | 1 | 415 | .184 | 0.01 |  |  |  |  |
|  |  |  | Age |  |  |  |  |  | -0.14 | 0.10 | -1.50 | .184 |
| PuL | Linear | F | Model | 7.42 | 1 | 429 | .024 | 0.02 |  |  |  |  |
|  |  |  | Age |  |  |  |  |  | -2.00x10^-4^ | 1.00x10^-4^ | -2.72 | .024 |
|  |  | M | Model | 0.02 | 1 | 418 | .900 | 5.47x10^-5^ |  |  |  |  |
|  |  |  | Age |  |  |  |  |  | -1.05x10^-5^ | 1.00x10^-4^ | -0.15 | .900 |
|  | Quadratic | F | Model | 3.71 | 2 | 428 | .060 | 0.02 |  |  |  |  |
|  |  |  | Age |  |  |  |  |  | -0.004 | 0.001 | -2.72 | .024 |
|  |  |  | Age^2^ |  |  |  |  |  | -0.001 | 0.002 | -0.32 | .882 |
|  |  | M | Model | 1.94 | 2 | 417 | .190 | 0.01 |  |  |  |  |
|  |  |  | Age |  |  |  |  |  | -3.00x10^-4^ | 0.002 | -0.22 | .860 |
|  |  |  | Age^2^ |  |  |  |  |  | -0.003 | 0.002 | -1.91 | .083 |
|  | Cubic | F | Model | 4.22 | 3 | 427 | .023 | 0.03 |  |  |  |  |
|  |  |  | Age |  |  |  |  |  | -0.01 | 0.003 | -3.12 | .009 |
|  |  |  | Age^2^ |  |  |  |  |  | -0.001 | 0.002 | -0.58 | .728 |
|  |  |  | Age^3^ |  |  |  |  |  | 0.004 | 0.002 | 2.17 | .070 |
|  |  | M | Model | 6.70 | 3 | 416 | 5.36x10^-4^ | 0.05 |  |  |  |  |
|  |  |  | Age |  |  |  |  |  | -0.02 | 0.004 | -3.91 | 3.31x10^-4^ |
|  |  |  | Age^2^ |  |  |  |  |  | -0.01 | 0.002 | -2.70 | .013 |
|  |  |  | Age^3^ |  |  |  |  |  | 0.01 | 0.002 | 4.01 | 2.30x10^-4^ |
|  | Log | F | Model | 5.69 | 1 | 429 | .047 | 0.01 |  |  |  |  |
|  |  |  | Age |  |  |  |  |  | -0.01 | 0.002 | -2.39 | .047 |
|  |  | M | Model | 0.57 | 1 | 418 | .501 | 0.001 |  |  |  |  |
|  |  |  | Age |  |  |  |  |  | 0.002 | 0.003 | 0.75 | .501 |
|  | S-curve | F | Model | 4.22 | 1 | 429 | .090 | 0.01 |  |  |  |  |
|  |  |  | Age |  |  |  |  |  | 0.16 | 0.08 | 2.05 | .090 |
|  |  | M | Model | 1.67 | 1 | 418 | .247 | 0.004 |  |  |  |  |
|  |  |  | Age |  |  |  |  |  | -0.12 | 0.09 | -1.29 | .247 |
| PuM | Linear | F | Model | 0.20 | 1 | 430 | .799 | 4.68X10^-4^ |  |  |  |  |
|  |  |  | Age |  |  |  |  |  | -3.34X10^-5^ | 1.00X10^-4^ | -0.45 | .799 |
|  |  | M | Model | 6.48 | 1 | 416 | .019 | 0.02 |  |  |  |  |
|  |  |  | Age |  |  |  |  |  | 2.00X10^-4^ | 1.00X10^-4^ | 2.55 | .019 |
|  | Quadratic | F | Model | 0.12 | 2 | 429 | .957 | 0.001 |  |  |  |  |
|  |  |  | Age |  |  |  |  |  | -0.001 | 0.002 | -0.44 | .802 |
|  |  |  | Age^2^ |  |  |  |  |  | -4.00X10^-4^ | 0.002 | -0.23 | .939 |
|  |  | M | Model | 5.66 | 2 | 415 | .007 | 0.03 |  |  |  |  |
|  |  |  | Age |  |  |  |  |  | 0.01 | 0.002 | 2.78 | .010 |
|  |  |  | Age^2^ |  |  |  |  |  | -0.004 | 0.002 | -2.19 | .045 |
|  | Cubic | F | Model | 0.09 | 3 | 428 | .981 | 0.001 |  |  |  |  |
|  |  |  | Age |  |  |  |  |  | -1.00X10^-4^ | 0.004 | -0.03 | .985 |
|  |  |  | Age^2^ |  |  |  |  |  | -4.00X10^-4^ | 0.002 | -0.19 | .951 |
|  |  |  | Age^3^ |  |  |  |  |  | -4.00X10^-4^ | 0.002 | -0.16 | .954 |
|  |  | M | Model | 4.32 | 3 | 414 | .009 | 0.03 |  |  |  |  |
|  |  |  | Age |  |  |  |  |  | -3.00X10^-4^ | 0.004 | -0.07 | .945 |
|  |  |  | Age^2^ |  |  |  |  |  | -0.01 | 0.002 | -2.44 | .024 |
|  |  |  | Age^3^ |  |  |  |  |  | 0.003 | 0.002 | 1.27 | .254 |
|  | Log | F | Model | 0.28 | 1 | 430 | .763 | 0.001 |  |  |  |  |
|  |  |  | Age |  |  |  |  |  | -0.002 | 0.003 | -0.53 | .763 |
|  |  | M | Model | 8.54 | 1 | 416 | .007 | 0.02 |  |  |  |  |
|  |  |  | Age |  |  |  |  |  | 0.01 | 0.003 | 2.92 | .007 |
|  | S-curve | F | Model | 0.88 | 1 | 430 | .464 | 0.002 |  |  |  |  |
|  |  |  | Age |  |  |  |  |  | 0.09 | 0.09 | 0.94 | .464 |
|  |  | M | Model | 8.55 | 1 | 416 | .007 | 0.02 |  |  |  |  |
|  |  |  | Age |  |  |  |  |  | -0.34 | 0.11 | -2.92 | .007 |
| VA | Linear | F | Model | 2.17 | 1 | 429 | .231 | 0.01 |  |  |  |  |
|  |  |  | Age |  |  |  |  |  | -2.00x10^-4^ | 1.00x10^-4^ | -1.47 | .231 |
|  |  | M | Model | 3.04 | 1 | 420 | .116 | 0.01 |  |  |  |  |
|  |  |  | Age |  |  |  |  |  | -2.00x10^-4^ | 1.00x10^-4^ | -1.74 | .116 |
|  | Quadratic | F | Model | 12.30 | 2 | 428 | 7.30x10^-5^ | 0.05 |  |  |  |  |
|  |  |  | Age |  |  |  |  |  | -0.003 | 0.002 | -1.58 | .191 |
|  |  |  | Age^2^ |  |  |  |  |  | -0.01 | 0.003 | -4.36 | 1.65x10^-4^ |
|  |  | M | Model | 2.81 | 2 | 419 | .089 | 0.01 |  |  |  |  |
|  |  |  | Age |  |  |  |  |  | -0.01 | 0.003 | -1.82 | .099 |
|  |  |  | Age^2^ |  |  |  |  |  | -0.004 | 0.003 | -1.40 | .212 |
|  | Cubic | F | Model | 9.63 | 3 | 427 | 4.75x10^-5^ | 0.06 |  |  |  |  |
|  |  |  | Age |  |  |  |  |  | -0.02 | 0.01 | -2.78 | .023 |
|  |  |  | Age^2^ |  |  |  |  |  | -0.01 | 0.003 | -4.84 | 3.36x10^-5^ |
|  |  |  | Age^3^ |  |  |  |  |  | 0.01 | 0.003 | 2.33 | .053 |
|  |  | M | Model | 5.89 | 3 | 418 | .002 | 0.04 |  |  |  |  |
|  |  |  | Age |  |  |  |  |  | -0.03 | 0.01 | -3.95 | 2.83x10^-4^ |
|  |  |  | Age^2^ |  |  |  |  |  | -0.01 | 0.003 | -2.08 | .057 |
|  |  |  | Age^3^ |  |  |  |  |  | 0.01 | 0.003 | 3.51 | .001 |
|  | Log | F | Model | 0.01 | 1 | 429 | .957 | 2.03x10^-5^ |  |  |  |  |
|  |  |  | Age |  |  |  |  |  | 4.00x10^-4^ | 0.004 | 0.09 | .957 |
|  |  | M | Model | 0.99 | 1 | 420 | .373 | 0.002 |  |  |  |  |
|  |  |  | Age |  |  |  |  |  | -0.01 | 0.01 | -1.00 | .373 |
|  | S-curve | F | Model | 2.59 | 1 | 429 | .186 | 0.01 |  |  |  |  |
|  |  |  | Age |  |  |  |  |  | -0.19 | 0.12 | -1.61 | .186 |
|  |  | M | Model | 0.01 | 1 | 420 | .941 | 1.55x10^-5^ |  |  |  |  |
|  |  |  | Age |  |  |  |  |  | 0.01 | 0.15 | 0.08 | .941 |
| VL | Linear | F | Model | 0.01 | 1 | 439 | .957 | 2.79x10^-5^ |  |  |  |  |
|  |  |  | Age |  |  |  |  |  | 6.80x10^-6^ | 1.00x10^-4^ | 0.11 | .957 |
|  |  | M | Model | 35.45 | 1 | 421 | 3.59x10^-8^ | 0.08 |  |  |  |  |
|  |  |  | Age |  |  |  |  |  | 4.00x10^-4^ | 1.00x10^-4^ | 5.95 | 3.59x10^-8^ |
|  | Quadratic | F | Model | 0.19 | 2 | 438 | .943 | 0.001 |  |  |  |  |
|  |  |  | Age |  |  |  |  |  | -0.001 | 0.001 | -0.39 | .841 |
|  |  |  | Age^2^ |  |  |  |  |  | -0.001 | 0.002 | -0.46 | .799 |
|  |  | M | Model | 23.49 | 2 | 420 | 1.62x10^-9^ | 0.10 |  |  |  |  |
|  |  |  | Age |  |  |  |  |  | 0.01 | 0.001 | 5.59 | 2.09x10^-7^ |
|  |  |  | Age^2^ |  |  |  |  |  | -0.003 | 0.002 | -2.07 | .057 |
|  | Cubic | F | Model | 3.33 | 3 | 437 | .052 | 0.02 |  |  |  |  |
|  |  |  | Age |  |  |  |  |  | -0.01 | 0.003 | -2.94 | .015 |
|  |  |  | Age^2^ |  |  |  |  |  | -0.002 | 0.002 | -1.06 | .397 |
|  |  |  | Age^3^ |  |  |  |  |  | 0.01 | 0.002 | 3.14 | .009 |
|  |  | M | Model | 20.21 | 3 | 419 | 4.52x10^-11^ | 0.13 |  |  |  |  |
|  |  |  | Age |  |  |  |  |  | 0.002 | 0.003 | 0.51 | .660 |
|  |  |  | Age^2^ |  |  |  |  |  | -0.002 | 0.002 | -1.48 | .186 |
|  |  |  | Age^3^ |  |  |  |  |  | 0.003 | 0.002 | 1.98 | .072 |
|  | Log | F | Model | 1.14 | 1 | 439 | .395 | 0.003 |  |  |  |  |
|  |  |  | Age |  |  |  |  |  | 0.002 | 0.002 | 1.07 | .395 |
|  |  | M | Model | 52.15 | 1 | 421 | 4.40x10^-11^ | 0.11 |  |  |  |  |
|  |  |  | Age |  |  |  |  |  | 0.02 | 0.002 | 7.22 | 4.40x10^-11^ |
|  | S-curve | F | Model | 4.03 | 1 | 439 | .097 | 0.01 |  |  |  |  |
|  |  |  | Age |  |  |  |  |  | -0.11 | 0.06 | -2.01 | .097 |
|  |  | M | Model | 55.55 | 1 | 421 | 1.58x10^-11^ | 0.12 |  |  |  |  |
|  |  |  | Age |  |  |  |  |  | -0.41 | 0.05 | -7.45 | 1.58x10^-11^ |
| VPL | Linear | F | Model | 5.27 | 1 | 429 | .056 | 0.01 |  |  |  |  |
|  |  |  | Age |  |  |  |  |  | -1.00x10^-4^ | 1.00x10^-4^ | -2.30 | .056 |
|  |  | M | Model | 30.93 | 1 | 418 | 2.35x10^-7^ | 0.07 |  |  |  |  |
|  |  |  | Age |  |  |  |  |  | 3.00x10^-4^ | 1.00x10^-4^ | 5.56 | 2.35x10^-7^ |
|  | Quadratic | F | Model | 3.21 | 2 | 428 | .090 | 0.02 |  |  |  |  |
|  |  |  | Age |  |  |  |  |  | -0.003 | 0.001 | -2.45 | .044 |
|  |  |  | Age^2^ |  |  |  |  |  | -0.001 | 0.001 | -0.90 | .481 |
|  |  | M | Model | 23.78 | 2 | 417 | 1.31x10^-9^ | 0.10 |  |  |  |  |
|  |  |  | Age |  |  |  |  |  | 0.01 | 0.001 | 5.73 | 1.07x10^-7^ |
|  |  |  | Age^2^ |  |  |  |  |  | -0.004 | 0.001 | -2.82 | .009 |
|  | Cubic | F | Model | 3.55 | 3 | 427 | .044 | 0.02 |  |  |  |  |
|  |  |  | Age |  |  |  |  |  | -0.01 | 0.003 | -2.80 | .022 |
|  |  |  | Age^2^ |  |  |  |  |  | -0.002 | 0.002 | -1.12 | .373 |
|  |  |  | Age^3^ |  |  |  |  |  | 0.003 | 0.002 | 1.96 | .103 |
|  |  | M | Model | 17.98 | 3 | 416 | 5.16x10^-10^ | 0.12 |  |  |  |  |
|  |  |  | Age |  |  |  |  |  | 4.00x10^-4^ | 0.003 | 0.14 | .901 |
|  |  |  | Age^2^ |  |  |  |  |  | -0.003 | 0.001 | -2.20 | .043 |
|  |  |  | Age^3^ |  |  |  |  |  | 0.0033 | 0.002 | 2.10 | .055 |
|  | Log | F | Model | 3.04 | 1 | 429 | .144 | 0.01 |  |  |  |  |
|  |  |  | Age |  |  |  |  |  | -0.004 | 0.002 | -1.74 | .144 |
|  |  | M | Model | 44.23 | 1 | 418 | 7.58x10^-10^ | 0.10 |  |  |  |  |
|  |  |  | Age |  |  |  |  |  | 0.01 | 0.002 | 6.65 | 7.58x10^-10^ |
|  | S-curve | F | Model | 1.58 | 1 | 429 | .309 | 0.004 |  |  |  |  |
|  |  |  | Age |  |  |  |  |  | 0.07 | 0.06 | 1.26 | .309 |
|  |  | M | Model | 45.27 | 1 | 418 | 5.16X10^-10^ | 0.010 |  |  |  |  |
|  |  |  | Age |  |  |  |  |  | -0.43 | 0.06 | -6.73 | 5.16X10^-10^ |

*Note.* Table includes models of five curves tested for each of the 14 thalamic nuclei within female and male participants separately.

Supplementary Table S4: AIC Comparisons for 14 Thalamic Nuclei

| Nuclei | Curve | Female Models | | Male Models | |
| --- | --- | --- | --- | --- | --- |
|  |  | Original | Weighted | Original | Weighted |
| AV | | | | | |
|  | Linear | - | n.s. | - | -1199.36 |
|  | Quadratic | - | **-1285.78** | - | -1210.58 |
|  | Cubic | - | n.s. | - | **-1218.09** |
|  | Log | - | -1275.02 | - | -1209.18 |
|  | S-Curve | - | -1213.28 | - | -1166.43 |
| IL | | | | | |
|  | Linear | - | n.s. | - | -1819.27 |
|  | Quadratic | - | n.s. | - | n.s. |
|  | Cubic | - | **-1859.51** | - | -1825.88 |
|  | Log | - | n.s. | - | **-1826.46** |
|  | S-Curve | - | n.s. | - | -1715.27 |
| LGN | | | | | |
|  | Linear | - | n.s. | - | -1749.02 |
|  | Quadratic | - | n.s. | - | **-1769.67** |
|  | Cubic | - | n.s. | - | n.s. |
|  | Log | - | **-1801.53** | - | -1762.48 |
|  | S-Curve | - | -1744.64 | - | -1717.56 |
| LP | | | | | |
|  | Linear | - | n.s. | - | n.s. |
|  | Quadratic | - | n.s. | - | -1254.69 |
|  | Cubic | - | n.s. | - | **-1256.88** |
|  | Log | - | n.s. | - | n.s. |
|  | S-Curve | - | n.s. | - | n.s. |
| MDl | | | | | |
|  | Linear | - | -1861.95 | - | n.s. |
|  | Quadratic | - | -1864.68 | - | -1790.33 |
|  | Cubic | - | **-1875.34** | - | **-1809.75** |
|  | Log | - | n.s. | - | -1798.99 |
|  | S-Curve | - | n.s. | - | -1656.66 |
| MDm | | | | | |
|  | Linear | - | -1818.17 | - | n.s. |
|  | Quadratic | - | -1813.44 | - | -1757.1 |
|  | Cubic | - | **-1828.89** | - | **-1774.32** |
|  | Log | - | n.s. | - | n.s. |
|  | S-Curve | - | n.s. | - | -1631.88 |
| MGN | | | | | |
|  | Linear | - | n.s. | - | n.s. |
|  | Quadratic | - | n.s. | - | **-1167.61** |
|  | Cubic | - | n.s. | - | n.s. |
|  | Log | - | n.s. | - | -1149.59 |
|  | S-Curve | - | n.s. | - | -1161.58 |
| PuA | | | | | |
|  | Linear | - | -1626.77 | -1545.37 | - |
|  | Quadratic | - | n.s. | **-1560.56** | - |
|  | Cubic | - | n.s. | n.s. | - |
|  | Log | - | **-1629.19** | -1555.8 | - |
|  | S-Curve | - | -1548.42 | -1480.17 | - |
| PuI | | | | | |
|  | Linear | -1850.21 | - | - | n.s. |
|  | Quadratic | n.s. | - | - | n.s. |
|  | Cubic | **-1859.9** | - | - | **-1728.94** |
|  | Log | n.s. | - | - | n.s. |
|  | S-Curve | n.s. | - | - | n.s. |
| PuL | | | | | |
|  | Linear | - | **-1869.49** | - | n.s. |
|  | Quadratic | - | n.s. | - | n.s. |
|  | Cubic | - | n.s. | - | **-1737.81** |
|  | Log | - | -1869.29 | - | n.s. |
|  | S-Curve | - | n.s. | - | n.s. |
| PuM | | | | | |
|  | Linear | - | n.s. | -1651.02 | - |
|  | Quadratic | - | n.s. | **-1653.81** | - |
|  | Cubic | - | n.s. | n.s. | - |
|  | Log | - | n.s. | -1653.06 | - |
|  | S-Curve | - | n.s. | -1538.24 | - |
| VA | | | | | |
|  | Linear | - | n.s. | - | n.s. |
|  | Quadratic | - | -1416.77 | - | n.s. |
|  | Cubic | - | n.s. | - | -1312.61 |
|  | Log | - | n.s. | - | n.s. |
|  | S-Curve | - | n.s. | - | n.s. |
| VL | | | | | |
|  | Linear | - | n.s. | - | -1870.35 |
|  | Quadratic | - | n.s. | - | n.s. |
|  | Cubic | - | n.s. | - | n.s. |
|  | Log | - | n.s. | - | **-1881.35** |
|  | S-Curve | - | n.s. | - | -1782.1 |
| VPL | | | | | |
|  | Linear | - | n.s. | - | -1888.73 |
|  | Quadratic | - | n.s. | - | -1899.58 |
|  | Cubic | - | n.s. | - | n.s. |
|  | Log | - | n.s. | - | **-1900.42** |
|  | S-Curve | - | n.s. | - | -1824.24 |

*Note.* Lowest AIC values are shown in bold font. n.s. = not significant either before or after multiple comparison correction.
